# Defining the roles of NKG7 expressed by CD4^+^ and CD8^+^ T cells during malaria

**DOI:** 10.1101/2025.09.11.675499

**Authors:** Teija C-M. Frame, Fabian de Labastida Rivera, Nicholas L. Dooley, Jessica A. Engel, Jinrui Na, Yulin Wang, Luzia Bukali, Susanna S. Ng, Patrick T. Bunn, Lynette Beattie, Timothy William, Matthew J. Grigg, Nicholas M. Anstey, James S. McCarthy, Bridget E. Barber, Michelle J. Boyle, Christian R. Engwerda

## Abstract

Malaria, caused by *Plasmodium* parasites, is a significant global health issue. CD4^+^ and CD8^+^ T cells are important for immunity against *Plasmodium* infections, but the specific roles of many immune-related effector molecules in T cells remain poorly defined. Here, we investigated the function of NK cell granule protein 7 (NKG7) in T cells during malaria, focusing on its role in CD4^+^ and CD8^+^ T cells in *Plasmodium* blood-stage responses. In a non-lethal malaria model, NKG7 played a protective role in CD4^+^ T cell responses, affecting pro-inflammatory T helper 1 (Th1), IL-10-producing type 1 regulatory (Tr1), and T follicular helper cell development. In a model of cerebral malaria, NKG7 was shown to have a cell-intrinsic role in CD4^+^ T cells for perforin and granzyme B expression, as well as the development of Tr1 cells. Human investigations involving peripheral blood mononuclear cells from volunteers participating in controlled human *P. falciparum* malaria infection studies, as well as endemic country patients with *P. falciparum* and *P. vivax* malaria, corroborated these findings. High NKG7 expression in T cells from *Plasmodium*-infected humans was observed, as well as differences in NKG7 expression based on the infecting *Plasmodium* species. NKG7 expression was associated with both cytotoxic and non-cytotoxic T cells, indicating varied functions following infection. These results advance our understanding about NKG7’s role in T cell-mediated malaria immunity and suggest potential for targeting NKG7 to improve outcomes following *Plasmodium* infection.

## Introduction

Malaria has a profound impact on human health, with 263 million cases causing an estimated 597,000 deaths in 83 endemic countries in 2023[1]. The COVID-19 pandemic intensified malaria incidence, leading to an estimated 13.4 million additional cases from 2019 to 2021[2], with recent global events further threatening the stability of malaria control programs. Africa bears the heaviest burden, accounting for 94% of malaria cases and 95% of malaria deaths, with children under five being the most affected[1]. *Plasmodium* parasites, particularly *P. falciparum*, and to a lesser extent *P. vivax*, cause most disease through complex interactions between parasites and human hosts[1, 3]. Animal models, such as rodent malaria infections, help advance understanding of disease pathogenesis and immunity[4–6], complementing *ex vivo* studies of clinical samples from malaria patients. Both animal and human studies have highlighted key roles for both CD4^+^ and CD8^+^ T cells in pathogenesis and immunity in malaria[3, 7]. An improved understanding about these cells will be needed for the development of second-generation malaria vaccines.

*Plasmodium*-specific CD4⁺ T cells are crucial mediators of protective immunity against malaria following natural exposure or vaccination[8–11], encompassing subsets such as T helper 1 (Th1), type I regulatory (Tr1), and T follicular helper (Tfh) cells that play important roles in both liver and blood stages of infection[3, 12]. Th1 cells, characterized by IFNγ production, are essential for controlling parasite burdens by activating macrophages and influencing B cell responses[13–18], though they can also contribute to malaria pathology by promoting atypical memory B cell development, inflammation and anaemia [19, 20]. Tr1 cells, which produce IL-10, help regulate inflammation but may hinder effective parasite control, with their presence linked to both higher parasitemia and reduced severe disease risk, highlighting their complex role in immune modulation[3, 12, 21–23]. Tfh cells are vital for fostering protective antibody responses by supporting the maturation of germinal centre (GC) B cells, leading to somatic hypermutation and the production of high affinity antibodies that target various *Plasmodium* lifecycle stages through mechanisms including antibody-dependent cell cytotoxicity and phagocytosis[24–29]. Overall, these CD4⁺ T cell subsets orchestrate a balance between effective immunity and immune regulation to manage *Plasmodium* infections.

Along with CD4⁺ T cells, *Plasmodium*-specific CD8⁺ T cells are detected in people with natural exposure to malaria[30], and after vaccination[31]; these show responses against sporozoite, liver, and blood-stage antigens[32, 33]. These cells can be important for liver-stage immunity, where they may prevent progression to symptomatic blood-stage infection[8, 34, 35]. Antigen-presenting cells in lymph nodes and the spleen prime CD8⁺ T cells[36–38], which employ both cytolytic- and cytokine-mediated mechanisms, including IFNγ, TNF, perforin, granzyme and granulysin production, to protect against liver-stage *Plasmodium*[39–41]. Liver-resident CD8⁺ T cells, generated by specific vaccination strategies in pre-clinical malaria models, form a protective niche in hepatic sinusoids[42–45]. While CD8⁺ T cells have limited roles in the blood stage of *P. falciparum* due to the lack of MHC class I on mature erythrocytes[46], responses against blood-stage antigens in *P. vivax* infection can eliminate infected reticulocytes[47]. However, in experimental cerebral malaria (ECM) in mice, CD8⁺ T cells may drive brain pathology through granzyme B, perforin, and IFNγ-mediated mechanisms[48, 49], targeting vascular endothelial cells cross-presenting parasite antigens[50–52].

Mechanisms of CD4^+^ and CD8^+^ T cell activation and differentiation in malaria are incompletely understood. We previously identified key roles for the natural killer cell protein 7 (NKG7) expression by CD8^+^ T cells for driving inflammation in ECM by facilitating CD107a translocation and cellular killing[53]. Initially identified as NK cell-specific in 1990 [54], the *NKG7* gene encodes a 148-amino acid protein with cytotoxic granule membrane localization in NK cells, which translocates to the plasma membrane upon cell activation[55–59]. NKG7 is also highly expressed in cytotoxic CD4⁺ and CD8⁺ T cells[60–64], correlating with Th1 cell polarization[65–67], and implicated in immune responses against infections like HIV, cytomegalovirus and COVID-19[60, 61, 68]. Its expression is also linked to cytotoxic functions in cancer and autoimmune disorders[69–76], making NKG7 a significant biomarker and potential target in transplant medicine, cancer, and immune-mediated inflammatory diseases. Expanding on our previous work[53, 77], here we investigate the role of CD4^+^ T cell NKG7 in non-lethal and lethal experimental malaria models, as well as NKG7 expression by CD4^+^ and CD8^+^ T cells in clinical samples from humans infected with *P. falciparum* and *P. vivax*.

## Results

### NKG7 has a protective role during non-lethal experimental malaria following *Plasmodium yoelii* 17XNL infection

We previously showed a key role for NKG7 expressed by CD8^+^ T cells in the pathogenesis of ECM caused by infection of C57BL/6 mice with *P. berghei* ANKA (*Pb*A)[53]. A key limitation of the ECM model is the acute and lethal nature of *Pb*A infection. Therefore, to investigate the role of NKG7 in parasite control and the development of immunity against malaria, an experimental mouse model of non-lethal, resolving malaria caused by *P. yoelii* (*Py*17XNL) infection was employed. We first infected *Nkg7*-deficient (B6^Δ*Nkg*^) and wild-type (WT) control C57BL/6 mice, and measured the progression of blood parasitemia through the course of infection (Figure 1A). Higher frequencies of parasitised red blood cells (pRBCs) were observed in *Nkg7-*deficient mice, compared to their WT counterparts (Figure 1B). To assess the overall parasite burden more comprehensively, the area under the curve (AUC) was calculated from blood parasitemia data points, revealing a significant increase in parasite burden among *Nkg7*-deficient mice (*P* < 0.05; Figure 1C). These results indicate an important role for NKG7 in the control of *Py*17XNL infection.

**Figure 1.**
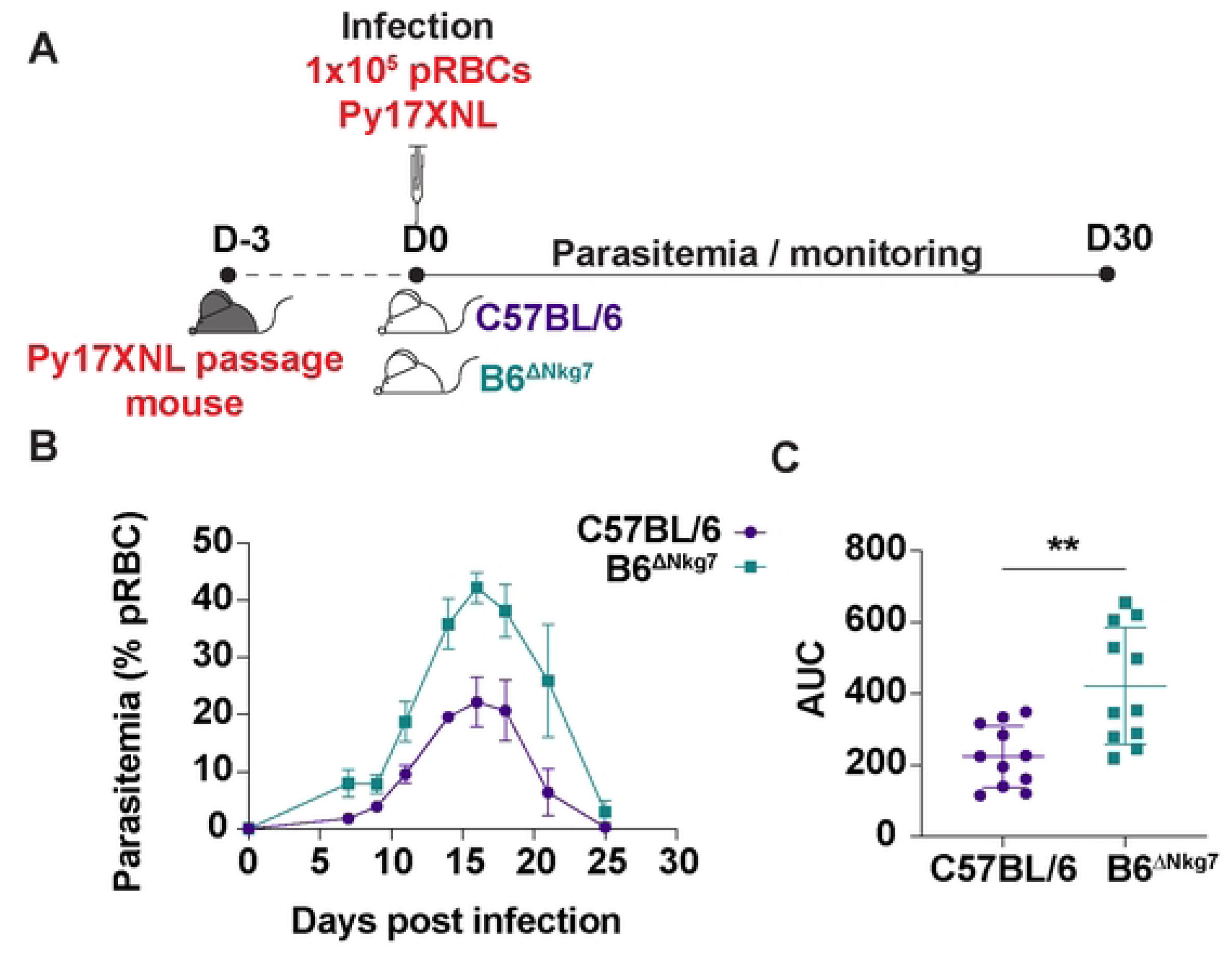
NKG7 has a protective role during non-lethal experimental malaria caused by *Plasmodium yoelii* 17XNL (*Py*17XNL) infection. C57BL/6 and B6.*Nkg7*^-/-^ (B6^Δ*Nkg7*^) mice were infected with non-lethal *Py*17XNL. Layout for the experimental timeline (A). Frequency of parasitised RBCs (pRBC) in peripheral blood of B6 and B6^Δ*Nkg7*^mice (B). Area Under the Curve (AUC), as a measure of parasite burden over the course of infection (C). Data is pooled from two independent experiments. A Mann-Whitney test was used to test for statistical significance, where ** P < 0.01. n = 11 per group.

### The role of NKG7 in Th1 and Tfh cell responses during non-lethal experimental malaria following *Plasmodium yoelii* 17XNL infection

To investigate whether NKG7 played a role in CD4^+^ T cell development in this malaria model, Th1 and Tfh cell responses were examined at two specific time points during infection. Both *Nkg7-*deficient mice and WT mice were infected with *Py*17XNL, and flow cytometry analysis was conducted on days 7 and 14 post-infection (p.i.) (Figure 2A & 2B). We found no significant differences in body weight, development of splenomegaly or increase in spleen cell numbers between *Nkg7*-deficient mice and WT mice at either time point during infection (Figure 2C). Upon treatment with PMA and ionomycin in the presence of monensin, significantly decreased frequencies of Th1 (IFNγ^+^ T-BET^+^) and Tr1 (IFNγ^+^ IL-10^+^) cells were observed in *Nkg7*-deficient mice at day 7 p.i., when compared to WT mice (*P* < 0.05; Figure 2D). Additionally, significantly reduced frequencies and numbers of Tfh (PD-1^+^ CXCR5^+^) cells were noted at day 14 p.i. in *Nkg7*-deficient mice compared to their WT counterparts (*P*< 0.05; Figure 2D). These data suggest a potential role for NKG7 in Th1 and Tr1 cell development during the first week of *Py*17XNL infection, as well as in Tfh cell development during the second week of *Py*17XNL infection.

**Figure 2.**
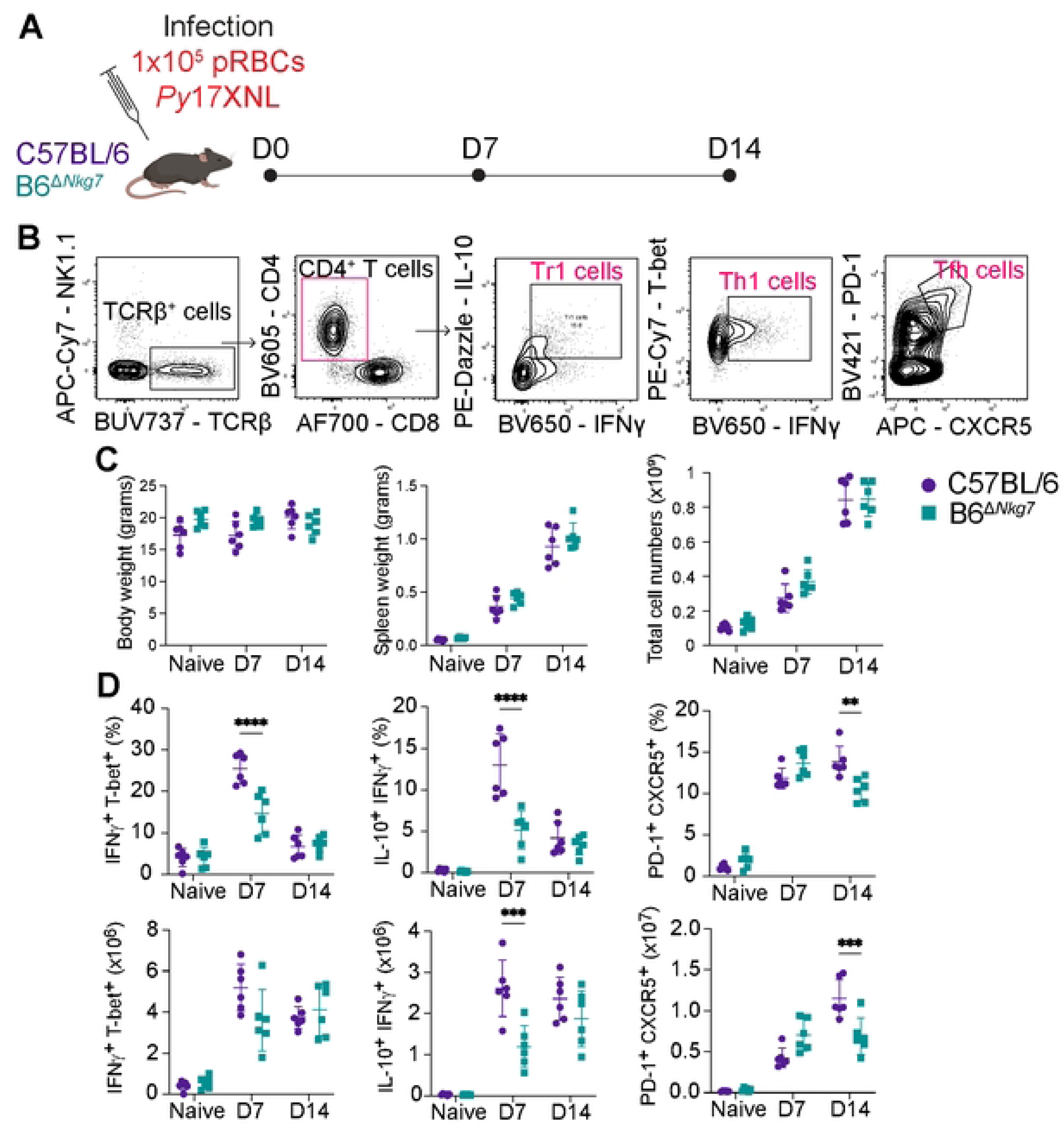
The role of NKG7 in CD4^+^ T cells during non-lethal experimental malaria caused by *Plasmodium yoelii* 17XNL (*Py*17XNL) infection. Schematic showing the experimental timeline (A). Gating strategy used to identify CD4^+^ T cell subsets (B). Body and spleen weights of naïve and *Py*17XNL infected C57BL/6 and B6.*Nkg7*^-/-^ (B6^Δ*Nkg7*^) mice. The right graph shows the total splenocyte numbers of naïve and infected C57BL/6 and B6^Δ*Nkg7*^ mice (C). Frequencies of IFNγ^+^ T-bet^+^ (Th1) cells and IFNγ^+^ IL-10^+^ (Tr1) cells as a percentage of TCRβ^+^ CD4^+^ T cells after stimulation with phorbol 12myristate 13-acetate and ionomycin. The corresponding numbers of these CD4^+^ T cell subsets in the spleen are also shown. The graphs on the right indicate frequencies of PD-1^+^ CXCR5^+^ (Tfh) cells as a frequency of TCRβ^+^ CD4^+^ T cells, and total number shown below (D). Data are from one representative experiment of two performed. A two-way ANOVA with Sidak’s multiple comparisons test was used to calculate statistical significance, where ** P < 0.01, *** P < 0.001, **** P < 0.0001. n = 6 mice per group.

These findings were supported when we examined the role of NKG7 in antigen-specific CD4^+^ T cells activation and differentiation in the lethal ECM model using PbT-II T cells. These are MHC-II-restricted TCR transgenic T cells which recognise a peptide derived from the *PbA* Hsp90 protein which is expressed by all rodent *Plasmodium* species[78]. To investigate cell-intrinsic roles for NKG7 in these cells, equal numbers of control (PbT-II^WT^) and *Nkg7*-deficient (PbT-II^Δ*Nkg7*^) cells were adoptively transferred into congenic *Ptprca* (*cd45.1*) recipient mice prior to *Pb*A infection. Cellular analysis was performed when the mice developed ECM at day 5 p.i. (Supplementary Figure 1A-B). After treatment with PMA and ionomycin, in the presence of monensin, despite a small, but significant increase in IFNγ^+^ PbT-II^Δ*Nkg7*^ cells, there was a decrease in Tr1 (IFNγ^+^ IL-10^+^) cell frequencies, compared with PbT-II^WT^ cells from the same animals (*P* < 0.05; Supplementary Figure 1C). PbT-II^Δ*Nkg7*^ cells also exhibited significantly lower frequencies of granzyme B-producing cells when compared to PbT-II^WT^ cells (*P* < 0.05), but no difference in CD107a expression (Supplementary Figure 1C), as previously observed for *Nkg7*-deficient CD8^+^ T cells[53]. We next examined their functional potential by assessing expression of different co-inhibitory receptors, but no differences in the expression of these molecules by PbT-II^WT^ and PbT-II^Δ*Nkg7*^ Tr1 cells was found (Supplementary Figure 1D). However, similar to granzyme B, a significant reduction in the frequency of perforin-producing PbT-II^Δ*Nkg7*^ cells was observed, when compared to the PbT-II^WT^ cells (*P* < 0.05; Supplementary Figure 1E). Together, these results identify roles for NKG7 in CD4^+^ T cells during *Pb*A infection in perforin and granzyme B expression, as well as the development of Tr1 cells.

### Increased *Nkg7* expression by Th1 and Tr1 cells during *Plasmodium yoelii* 17XNL infection

To characterise the cellular expression of NKG7 during *Py*17XNL infection, a *Nkg7*-transcriptional reporter mouse was utilised (Figure 3A). In these mice, cells in which *Nkg7* transcription is initiated switch from expressing TdTomato (red) to GFP (green)[53]. In the naïve state, the majority of NK cells expressed GFP, but this expression significantly decreased following infection (*P* < 0.05), as previously reported in experimental visceral leishmaniasis[53]. Conversely, the frequency of GFP^+^ cells increased significantly in NKT, CD4^+^ and CD8^+^ T cell subsets during the infection (*P* < 0.05; Figure 3B). The frequency of GFP-expressing CD4^+^ T cells doubled from the naïve state to day 7 p.i. and was sustained at day 14 p.i. (Figure 3B). To further investigate this increase, we next examined *Nkg7* expression within distinct CD4^+^ T cell subsets. Following activation with PMA and ionomycin, Th1 cells dominated in the naïve state, while Tr1 cells became dominant at days 7 and 14 p.i. (Figure 3C). The compartment representing other CD4^+^ T cells included cells that were neither Th1 cells, Tr1 cells, nor Treg cells. When analysing T cell subsets directly *ex vivo*, without further stimulation, a similar trend emerged with significantly increasing frequencies of GFP^+^ cells during the infection in CD4^+^ and CD8^+^ T cells (*P* < 0.05; Figure 3D), as well as in the CD4^+^ T cell subsets, including Th0 cells (T-BET^-^ CXCR6^-^), Th1 cells (T-BET^+^ CXCR6^+^), and other CD4^+^ T cells (non-Tfh T-BET^+^ CXCR6^+^) (*P* < 0.05; Figure 3E). We were unable to detect IL-10-producing Tr1 cells without *ex vivo* activation with mitogens in this analysis. Interestingly, the frequency of GFP^+^ Tfh cells (PD-1^+^ CXCR5^+^) remained consistent between the naïve state and both infection time points (Figure 3E). These data suggest that NKG7 is expressed across multiple CD4^+^ T cell subsets with enrichment in Th1 and Tr1 cells following *Py*17XNL infection.

**Figure 3.**
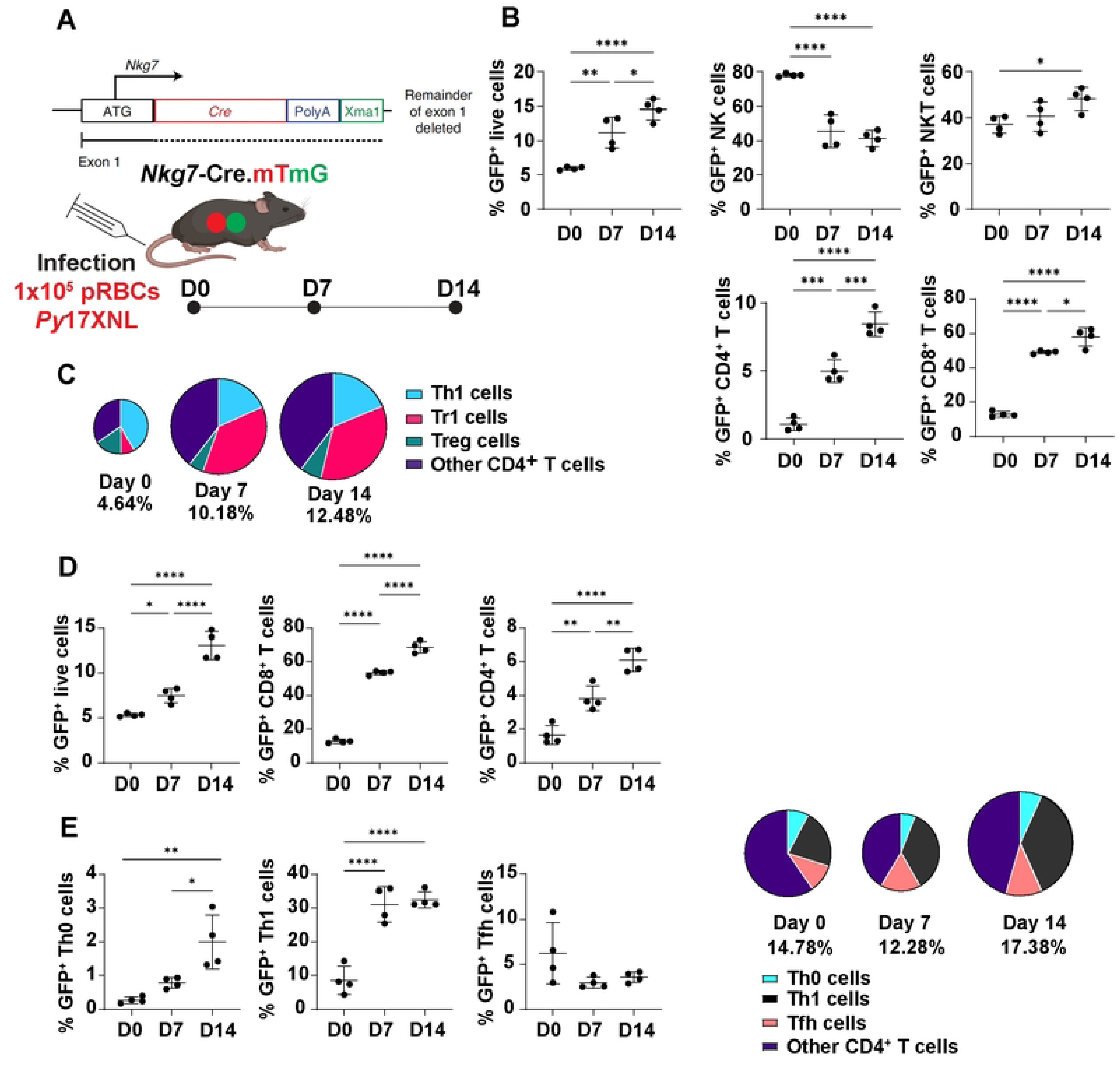
*Nkg7* expression increases in Th1 and Tr1 cells during non-lethal *Plasmodium yoelii* 17XNL (*Py*17XNL) infection. Schematic showing the experimental timeline for B6.*Nkg7*-reporter mice (*Nkg7*-cre x mT/mG) were infected with *Py*17XNL (A). Frequencies of GFP^+^ (*Nkg7*^+^) in lymphoid cell subsets, as indicated, during *Py*17XNL infection (B). Frequencies of GFP^+^ (*Nkg7*^+^) CD4^+^ IFNγ^+^ T-bet^+^ (Th1) cells, IFNγ^+^ IL-10^+^ (Tr1) cells, Foxp3^+^ regulatory T (Treg) cells and the remaining CD4^+^ T cells after stimulation with phorbal 12myristate 13-acetate and ionomycin, shown as a pie chart. The size of the compartments within the pie chart indicates the percentage of the CD4^+^ T cell subsets specified, within the GFP^+^ CD4^+^ T cell population (C). Frequencies of all GFP^+^ (*Nkg7*^+^) cells, CD8^+^ T cells and CD4^+^ T cells during *Py*17XNL infection directly *ex vivo* (D). CD4^+^ T cells were further divided into CXCR6^-^ T-bet^-^ (Th0) cells, CXCR6^+^ T-bet^+^ (Th1) cells, PD-1^+^ CXCR5^+^ (Tfh) cells and the remaining CD4^+^ T cells, shown as a pie chart. The size of the compartments within the pie chart indicates the percentage of the CD4^+^ T cell subsets specified, within the GFP^+^ CD4^+^ T cell population (E). Statistical significance was assessed by ordinary one-way ANOVA with Tukey’s multiple comparisons test, where * P < 0.05, ** P < 0.01, *** P < 0.001, **** P < 0.0001. n = 4 naïve or infected mice. Data are from one representative experiment of two performed.

### A cell intrinsic role for NKG7 in the development of Th1 and Tfh cells during *Plasmodium yoelii* 17XNL infection

To investigate whether NKG7 played a cell-intrinsic role in CD4^+^ T cells in the non-lethal malaria model, equal numbers of PbT-II^WT^ and PbT-II^Δ*Nkg7*^ cells were adoptively transferred into congenic Ptprca (*cd45.1*) recipient mice prior to *Py*17XNL infection. Cellular analyses were performed at both day 7 and day 14 p.i. to evaluate Th1, Tr1 and Tfh cell responses (Figure 4A & 4B). At day 7 p.i., following treatment with PMA and ionomycin in the presence of monensin to assess cellular potential, no discernible differences were observed in the frequencies of Th1 cells (IFNγ^+^ T-BET^+^) between the two PbT-II populations (Figure 4C). However, we observed a significant reduction in the frequencies of PbT-II^Δ*Nkg7*^ Tr1 cells (IFNγ^+^ IL-10^+^) (*P* < 0.05; Figure 4C). In unstimulated cells analysed *ex vivo*, a significant decrease was noted in PbT-II^Δ*Nkg7*^ Tfh cell (PD-1^+^ CXCR5^+^) frequencies, as well as PbT-II^Δ*Nkg7*^ Th1 (T-BET^+^ CXCR6^+^) cells (*P* < 0.05; Figure 4C). At day 14 p.i., in unstimulated cells, significant differences were only observed in Th1 (T-BET^+^ CXCR6^+^) cells (*P* < 0.05; Figure 4D). Taken together, these data provide support for a cell-intrinsic role of NKG7 in the development of Th1, Tr1 and Tfh cells at day 7 p.i. and Th1 cells at day 14 p.i..

**Figure 4.**
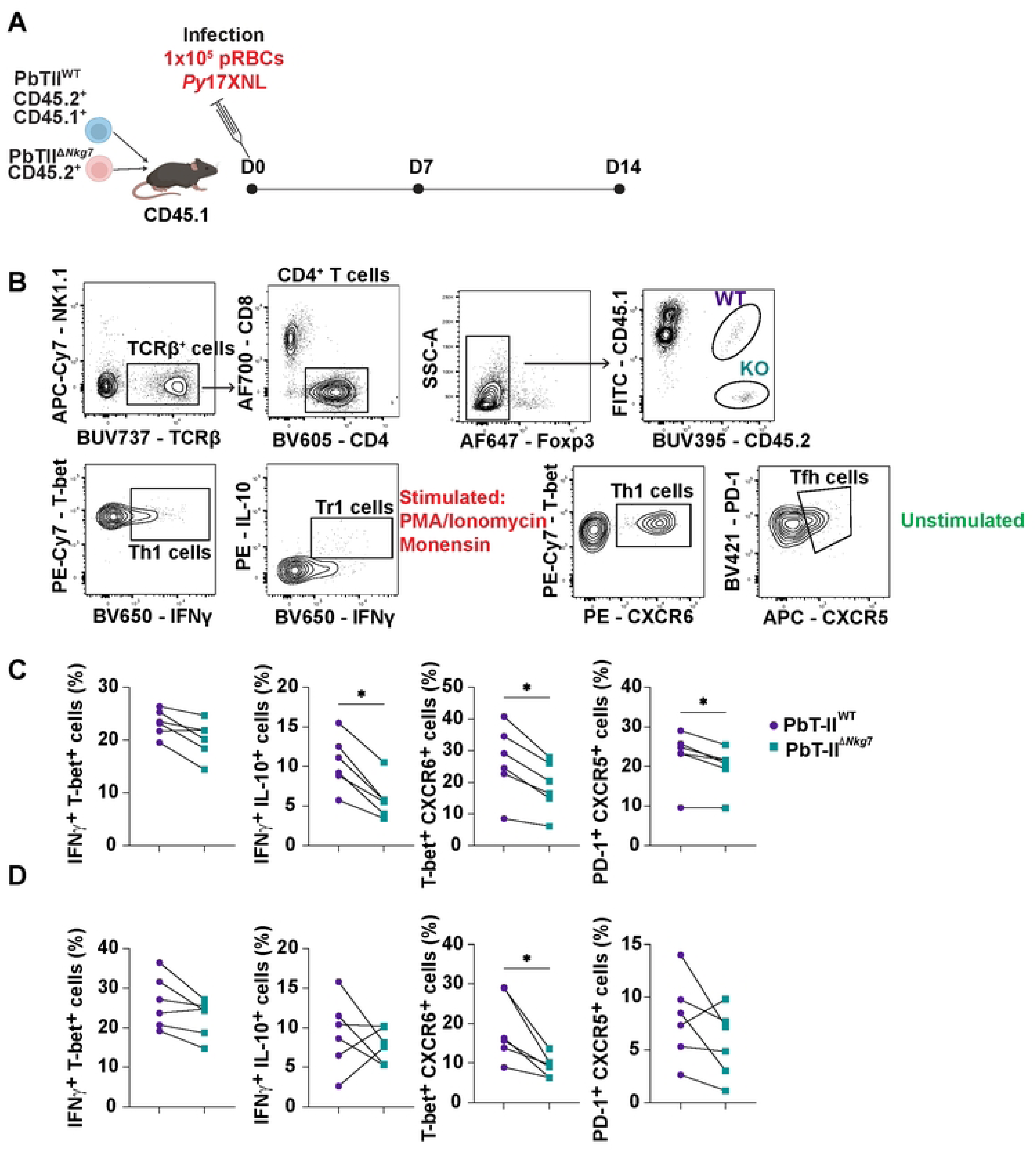
NKG7 has a cell intrinsic role in CD4^+^ T cell responses during non-lethal experimental malaria caused by *Plasmodium yoelii* 17XNL (*Py*17XNL) infection. Experimental timeline for co-transfer of transgenic PbT-II^WT^ and PbT-II^Δ*Nkg*7^ cells into B6.CD45.1 mice prior to infection with *Py*17XNL (A). Gating strategy used to identify and phenotype PbT-II cell subsets (B). Frequencies of co-transferred transgenic PbT-II^WT^ and PbT-II^Δ*Nkg*7^ IFNγ^+^ T-bet^+^ (Th1) cells and IFNγ^+^ IL-10^+^ (Tr1) cells as a proportion of TCRβ^+^ CD4^+^ T cells from the spleen at day 7 post infection after stimulation with phorbol ester and ionophore (two left panels). Frequencies of co-transferred transgenic PbT-II^WT^ and PbT-II^Δ*Nkg*7^ T-bet^+^ CXCR6^+^ cells and PD-1^+^ CXCR5^+^ (Tfh) cells as a proportion of TCRβ^+^ CD4^+^ T cells from the spleen at day 7 post infection *ex vivo* (no further stimulation; two right panels) (C). Frequency of the same PbT-II cells as above at day 14 post infection (E). A Wilcoxon test was used to define statistical significance, where * P < 0.05, n = 6 mice per group. Data are from one representative experiment of two performed.

### The expression of NKG7 during controlled human infection studies with *Plasmodium falciparum*

We next examined the pattern of NKG7 expression by human immune cells during malaria. To measure NKG7 expression in human peripheral blood mononuclear cells (PBMCs), we first tested the specificity of a previously reported monoclonal antibody[58]. We employed HEK293 cells transfected with human *NKG7-GFP* and the NK cell line NK92, for which CRISPR/cas9 was employed to delete *NKG7*. We readily detected NKG7 in *NKG7*-transfected HEK293 cells, but not in control cells, as well as NK92 wild-type cells, but not cells depleted of *NKG7*, where expression was significantly diminished at both mRNA and protein levels (Supplementary Figure 2). Thus, this anti-NKG7 antibody was included in flow cytometry antibody panels for further investigations.

We next examined blood from healthy, malaria-naïve volunteers participating in controlled human malaria infection (CHMI) studies with blood-stage *P. falciparum*[79]. This involved samples from 8 individuals (5/8 male, aged median 22, range 18-29); all were treated with an anti-parasitic drug 8-9 days after infection, when blood parasite level exceeded 5000 parasites/mL in most participants[80]. PBMCs were isolated from blood and examined before infection (day 0), at pre-treatment (day 8 p.i., prior to drug administration), after anti-parasitic treatment (day 15 p.i.) and at the end of the study (EOS; day 45 ± 2) (Figure 5A). Unsupervised dimensionality reduction using UMAP, followed by clustering of flow cytometry data, identified 30 distinct cell clusters, (Figure 5B). Cell clusters with high NKG7 expression also had high expression of the cytotoxic molecules GZMB, CD107a and KLRG1 (Figure 5C). We identified one CD4^+^ T cell cluster (C04) and five CD8^+^ T cell clusters (C07, C13, C14, C15 and C17) that had high expression of NKG7 (Figure 5D-G). The NKG7^+^ CD4^+^ T cell cluster (C04) displayed characteristics of an effector memory CD4^+^ T cell cluster (CD25^-^ CD127^+^ CCR7^-^) that expressed multiple activation markers, including CD49b, PD1, CD38, KLRG1, GZMB and CD107a (Figure 5D & F). The five NKG7^+^ CD8^+^ T cell clusters lacked CCR7 expression, and were identified as likely effector memory T cell clusters with varying levels of CD45RA expression. Three of the NKG7^+^ CD8^+^ T cell clusters (C07, C13 and C14) showed a similar cytotoxic phenotype with different combinations of KLRG1, CD38, and PD1 expression. Cluster 17 appeared to be a less differentiated CD8^+^ T cell cluster as it lacked expression of cytotoxic, activation or exhaustion markers and had low expression of KLRG1 and CCR7. Cluster 15 showed differentiation from the other four NKG7^+^ CD8^+^ T cell clusters and expressed high levels of CCR6, CD56 and KLRG1 and lacked expression of cytotoxic molecules (Figure 5D & G), suggesting it may represent either innate-like CD8^+^ T cells[81] or non-cytotoxic memory-like CD8^+^ T cells[82]. Examination of the differential abundance of cell clusters after infection (day 0 to day 8, day 0 to day 15, day 0 to EOS), revealed a significant increase (*P* < 0.05) in the abundance of the one NKG7^+^ CD4^+^ T cell cluster (C04) and one of the NKG7^+^ CD8^+^ T cell clusters (C07) when day 0 (pre-infection) was compared to day 15 (15 days p.i., 7 days post-drug treatment) (Figure 5E).

**Figure 5.**
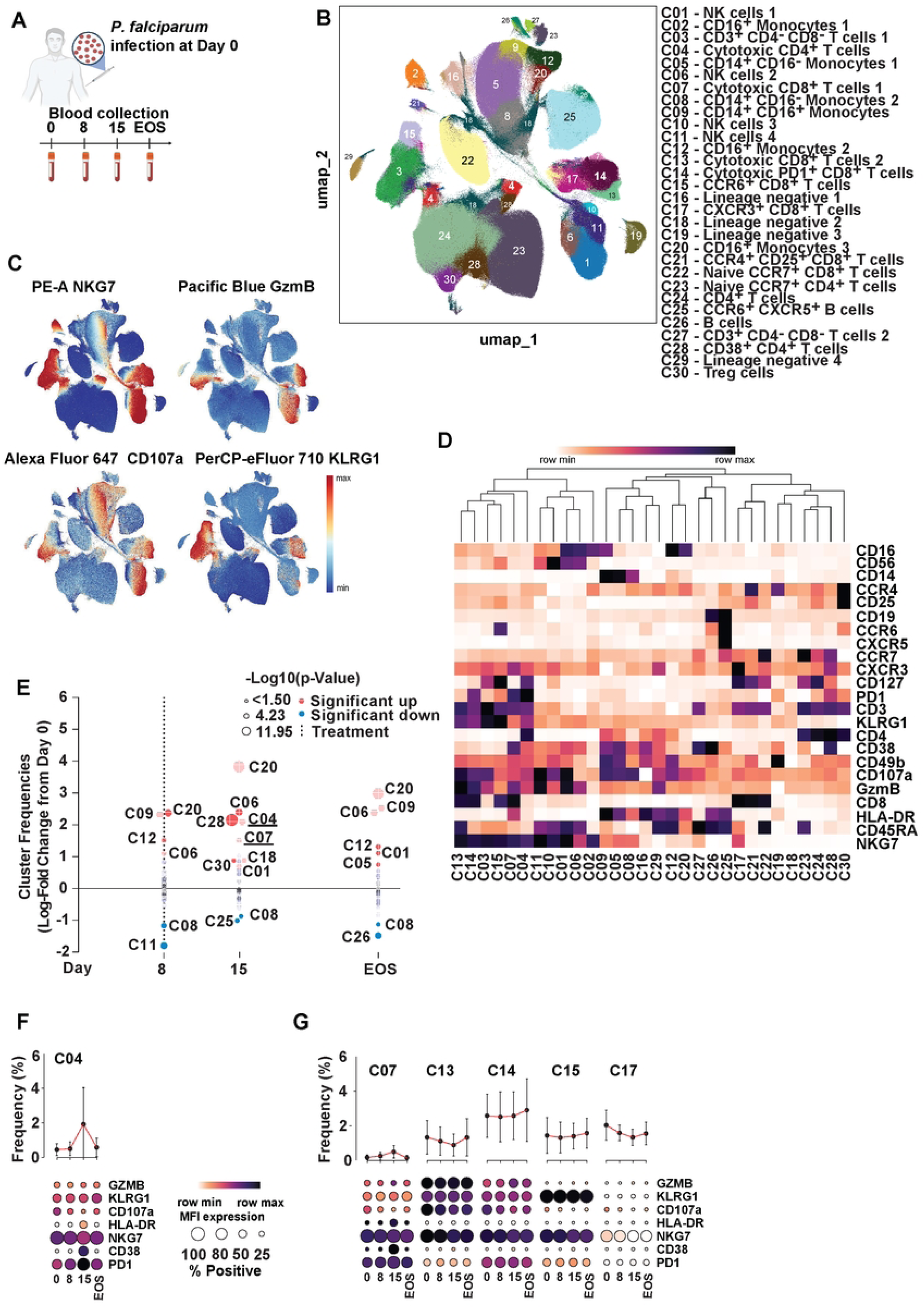
NKG7 expression correlated with cytotoxic and activated cells during controlled human malaria infection (CHMI). Schematic showing the experimental design for CHMI studies; malaria-naïve subjects were infected with blood-stage *Plasmodium falciparum* 3D7 followed by treatment with an anti-parasitic trial drug, 8-9 days after infection. Peripheral blood mononuclear cells (PBMCs) were isolated from blood samples that were collected for analysis before infection (day 0), pre-treatment (day 8 post infection (p.i.), prior to drug administration), post anti-parasitic treatment (day 15 p.i.) and at the end of the study (EOS) (A). Cryopreserved PBMCs were thawed and immunophenotyped by spectral flow cytometry. Post-acquisition analysis was conducted using OMIQ software for dimensionality reduction using unsupervised FLOWSOM clustering that yielded 30 clusters (B). Mean expression of cytotoxic markers NKG7, granzyme B (GzmB), CD107a, and KLRG1 are also shown separately on the UMAP (C). Expression of cell phenotyping markers were measured, and mean fluorescence intensity (MFI) of staining presented as a heatmap. n = 31 volunteer samples, day 0 (n = 8), day 8 (n = 8), day 15 (n=8), EOS (n = 7) (D). PBMCs were divided into 30 clusters and clusters that were significantly different in abundance between day 0 and 8, day 0 and 15, and day 0 and end of study (EOS) are indicated with red circles for significant increase and blue for significant decrease (significance assessed using EdgeR). T cell clusters of interest are underlined (E). Frequency (top) and Dot plots (bottom) depicting Cluster change, expression patterns and frequency of positive events for cytotoxic and activation markers in a CD4^+^ T cell cluster (F) and CD8^+^ T cell clusters (G) for the duration of the CHMI trial.

Further analysis of samples at day 15 p.i., when T cell activation appeared to peak, revealed significantly increased expression of CD38, PD1 and HLA-DR within the NKG7^+^ CD4^+^ T cell cluster (C04) (*P* < 0.05; Figure 5F & Supplementary Figure 3B). However, significantly decreased NKG7 expression from day 0 to day 15 was observed within this population, mirroring a decrease in CD107a expression (*P* < 0.05), and supporting a role for NKG7 in degranulation of effector molecules (Supplementary Figure 3B). Cluster 7, which was the only CD8^+^ T cell cluster that was significantly different in abundance from day 0 to day 15 p.i. (Figure 5E), had relatively high expression of CD38, HLA-DR and PD-1 (Figure 5G), suggesting a highly activated and exhausted cluster. Similar to the NKG7^+^ CD4^+^ T cell cluster C04, a significantly decreased expression of CD107a from day 0 to day 15 was also observed in all NKG7^+^ CD8^+^ T cell clusters (*P* < 0.05; Figure 5G & Supplementary Figure 3B). Together, these data provide new insights into CD4^+^ and CD8^+^ NKG7-expressing T cells during CHMI with multiple T cell subsets identified, each with a different functional potential.

### Species-specific differences in NKG7-expressing T cells during uncomplicated *Plasmodium falciparum* and *P. vivax* malaria

To test whether the NKG7 expression patterns observed in CHMI samples were also found during naturally acquired infection, we next investigated blood samples from patients admitted to a tertiary-referral hospital in Sabah, Malaysia, during 2010–2012, with uncomplicated *P. falciparum* or *P. vivax* malaria. Blood was collected at the time of presentation and one month post anti-parasitic treatment. These samples were classified into three groups: acute *P. falciparum* (*Pf*) infection (n=5), acute *P. vivax* (*Pv*) infection (n=6) and convalescence. The convalescence group included samples from 2 *Pf* and 6 *Pv* patients. Following analysis of flow cytometry data, 20 clusters were identified (Figure 6A), with the majority of NKG7-expressing clusters also expressing the cytotoxic molecules granulysin (GNLY) and/or granzyme B (GZMB; Figure 6B). Of the NKG7^+^ clusters, two CD4^+^ T cell clusters and four CD8^+^ T cell clusters were identified (Figure 6B & C). The two NKG7^+^ CD4^+^ T cell clusters (C05 and C20) appeared to be effector memory cell clusters as both had lost CD27 expression, with cluster 20 having high CD57 expression (Figure 6C). Three of the NKG7^+^ CD8^+^ T cell clusters were grouped together in the UMAP (C11, C17 and C18), with cluster 11 and cluster 17 overlapping, suggesting phenotypic and possible functional similarities (Figure 6A). Cluster 13 was clearly separated from the other three NKG7^+^ CD8^+^ T cells clusters in the UMAP (Figure 6A), and had high CD161 expression (Figure 6B & C), overlapping several NKG7^+^ CD8^+^ T cells clusters observed in CHMI samples (Figure 5).

**Figure 6.**
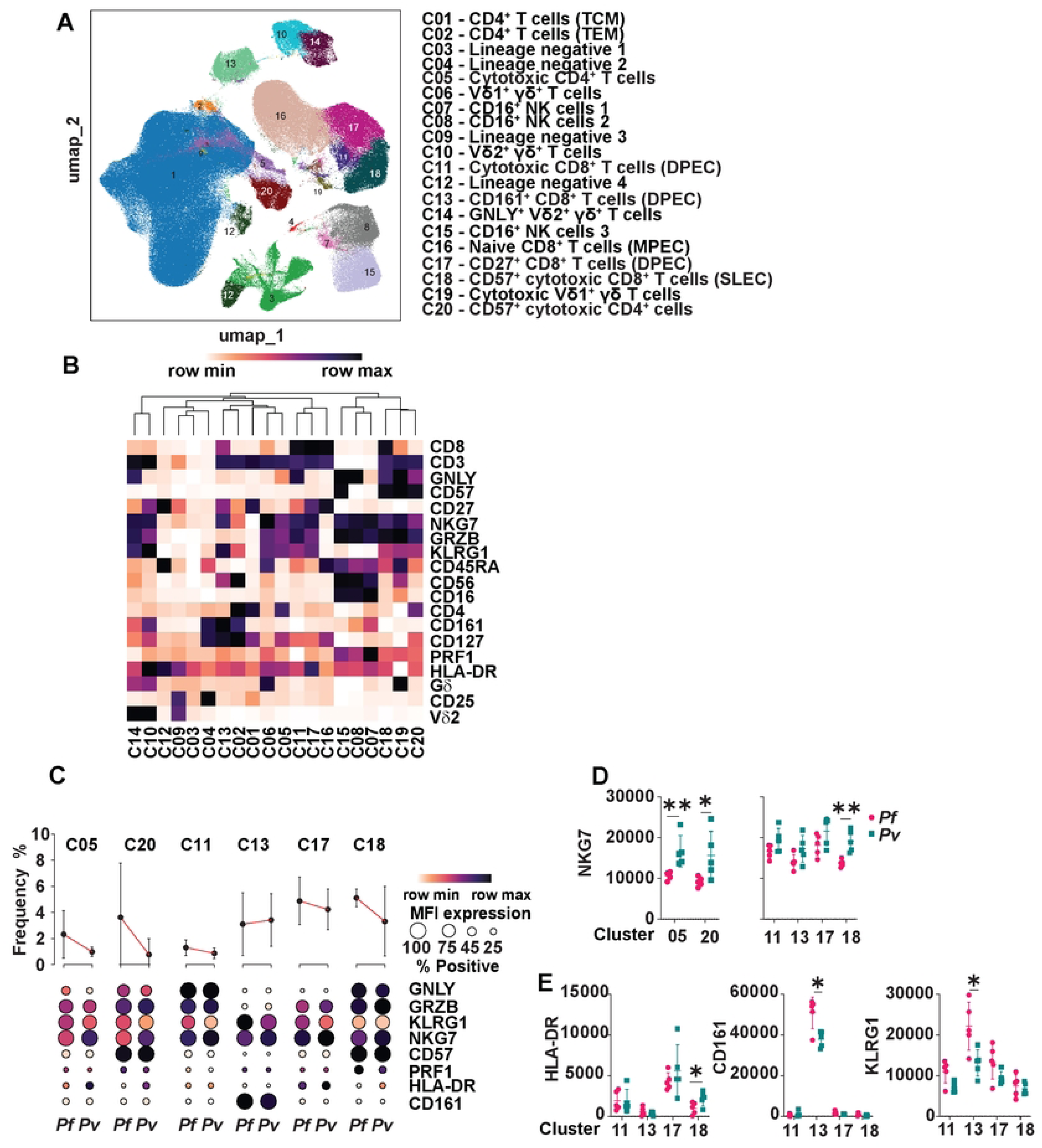
Differences in NKG7 expression during *P. falciparum* and *P. vivax* naturally-acquired malaria. Peripheral blood mononuclear cells (PBMCs) were isolated from blood samples collected from patients with naturally-acquired, uncomplicated malaria at the Queen Elizabeth Hospital in Sabah, acute *P. falciparum* (*Pf*; n = 5) and acute *P. vivax* (*Pv*; n =6). Post-treatment samples were also collected (n = 8). Cryopreserved PBMCs were thawed and immunophenotyped by spectral flow cytometry. Post-acquisition analysis was conducted using OMIQ software for dimensionality reduction using and unsupervised FLOWSOM clustering (A). Expression of cell protein markers was measured, and mean fluorescent intensity of staining presented as a heatmap (B). Samples were divided into 20 clusters and the frequency of CD4^+^ and CD8^+^ T cell clusters are shown (top) and dot plots (bottom) depicting Cluster change, expression patterns and frequency of positive events for cytotoxic and activation markers compared between *Pf* and *Pv* (C). NKG7 expression for CD4^+^ and CD8^+^ T cell clusters (D), and HLA-DR, CD162 and KLRG1 expression for all CD8^+^ T cell clusters are shown (E). Significance was assessed by ordinary one-way ANOVA with Tukey’s multiple comparisons test, where * P < 0.05, n = 5-6 patient samples per group.

By using the same analysis methods employed for CHMI samples, the two NKG7^+^ CD4^+^ T cell clusters (C05 and C20) were found to be the only clusters with significant differences in abundance when comparing the acute infections of the two different *Plasmodium* species, with both clusters increased during acute *Pf* infection relative to *Pv* (Supplementary Figure 4). When acute *Pf* infection was compared to convalescence, none of the four significantly different clusters included CD4^+^ or CD8^+^ T cells (Supplementary Figure 4), although two NKG7-expressing NK cell clusters were increased. However, one of the four clusters identified as being significantly increased in frequency in acute *Pv* infection, compared to convalescence, was cluster 11, a NKG7^+^ CD8^+^ T cell cluster, consistent with a role for cytotoxic CD8^+^ T cells in immune responses to *Pv* malaria[47] (Supplementary Figure 4).

We also compared differences within the NKG7^+^ CD4^+^ and CD8^+^ T cells clusters between patient groups. In both NKG7^+^ CD4^+^ T cell clusters, there was significantly higher expression of NKG7 expression in acute *Pv* infection, when compared to acute *Pf* infection (*P* < 0.05; Figure 6D). The only other difference between the NKG7^+^ CD4^+^ T cell clusters was CD57 expression, which was much higher overall in cluster 20, although no difference was observed between patient groups. This cluster also had a small increase in the overall expression of cytotoxic molecules (Figure 6C). The four NKG7^+^ CD8^+^ T cell clusters also had increased NKG7 expression during acute *Pv* infection when compared to acute *Pf* infection, but was only significantly increased in cluster 18 (*P* < 0.01; Figure 6D). Cluster 18 also had significantly higher expression of HLA-DR during *Pv* infection when compared to *Pf* infection (*P* < 0.05; Figure 6E) and overall had higher CD57 expression in both groups when compared to the other clusters. Cluster 13 was a non-cytotoxic cluster and had high levels of CD161 expression, which was significantly increased in acute *Pf* infection, when compared to acute *Pv* infection (*P* < 0.05; Figure 6E). KLRG1 expression in cluster 13 followed the same pattern when the two acute infections were compared (*P* < 0.05; Figure 6E). Overall, this data confirms that the NKG7 expression patterns found in CD4^+^ and CD8^+^ T cells in CHMI samples were similar to those observed in natural infection, with some notable differences between patients infected with different *Plasmodium* species.

## Discussion

CD4^+^ and CD8^+^ T cells play pivotal roles in host immune responses during malaria. They are essential for the killing and clearing of *Plasmodium* parasites and for the development of immunity[3, 83]. Recent research has shed light on the involvement of NKG7 in parasitic-causing diseases, including both leishmaniasis and malaria[53]. The study presented here was designed to elucidate the role of NKG7 in CD4^+^ and CD8^+^ T cells during malaria. Further to its known role in facilitating CD8^+^ T cell exocytosis and enhancing the cytotoxic function of CD8^+^ T cells during *Pb*A infection, we also found NKG7 was critical for development of IL-10-producing Tr1 cells and cytotoxic CD4^+^ T cells in this model. Additionally, NKG7 was shown to be associated with protection during non-lethal *Py*17XNL infection, and to contribute to the development of CD4^+^ T helper subsets, including Th1, Tr1, Tfh and cytotoxic CD4^+^ T cells. In human PBMCs obtained from individuals infected with either *P. falciparum* or *P. vivax*, NKG7 was found within cytotoxic cell subsets, and notably, within some distinctive, non-cytotoxic cell populations. Furthermore, we discovered NKG7 species-specific differences in samples from patients with *P. falciparum* and *P. vivax* malaria from Sabah, Malaysia. These findings contribute to our understanding about how the host’s immune system responds to malaria, and highlight the dual potential of targeting NKG7 activity for both stimulating immune responses against intracellular pathogens and applications for dampening inflammation.

The functions of CD8^+^ T cells have been extensively documented in *Plasmodium* infection, particularly regarding their role in mediating protection at the liver stage of disease[83, 84]. In agreement with numerous transcriptional studies implicating NKG7 in the function of cytotoxic effector CD8^+^ T cells[63, 64, 85–88], we have reported a key role for NKG7 in CD8^+^ T cell exocytosis and cytotoxicity. However, this was associated with NKG7-exacerbated parasite accumulation in tissues and the onset of ECM following *Pb*A infection[53]. Thus, *Nkg7*-deficient CD8^+^ T cells exhibit reduced expression of key cytotoxic molecules, such as granzyme B, perforin and CD107a, and a diminished capacity to eliminate target cells. Recent studies in other disease models have confirmed these findings for NKG7, emphasising a decreased killing ability and impaired degranulation in the absence of NKG7[63, 77, 89–92]. Of particular interest, in one of these studies[89], structural similarities were observed between the four membrane-spanning domains of NKG7 and the γ subunit of an L-type voltage-gated calcium channel. In this context, the authors were able to demonstrate that reduced NKG7 expression compromised calcium flux in CD8^+^ T cells[89]. This aligns with the hypothesis that NKG7 is involved in regulating important intracellular processes, such as intracellular calcium flux, which are crucial for numerous T cell functions. Together, these findings highlight the potential benefits of increasing NKG7 expression for enhancing cytotoxicity anti-parasitic responses. If NKG7 indeed improves the ability of CD8^+^ T cells to eliminate target cells, then the overexpression of this protein could represent a novel therapeutic strategy with broad applicability in various clinical contexts, including infectious diseases caused by intracellular pathogens, as well as cancer. In fact, a recent study demonstrated gain-of-function (tumour killing) in CAR T cells ectopically over-expressing NKG7 in pre-clinical cancer models[93]. However, the consequences of this on the outcome of immunotherapy, and other approaches to modulate NKG7 would have to be carefully monitored.

Th cell subsets play pivotal roles in the immune response to malaria. Th1 cells are essential for cell-mediated, anti-parasitic responses, while Tfh cells are critical for the development of antigen-specific B cells which produce antibodies required for immunity to malaria[3, 12, 83, 94]. The transcription factor T-BET, a master transcription factor for Th1 cell development, binds to the *NKG7* promoter, increasing the expression of *NKG7*, *PRF1* and *IFNG*[65]. NKG7 has also been implicated in other Th cell subsets, including recent reports of expression by cytotoxic CD4^+^ T cells[60–62, 95]. In experimental visceral leishmaniasis, NKG7 was found to be crucial for the activation of STAT4 and Th1 cell development[53]. Here, a protective role for NKG7 during *Py*17XNL infection was shown, as well as its involvement in the development and functions of CD4^+^ T cell subsets, including Th1, Tr1 and Tfh cells. NKG7 expression within CD4^+^ T cell subsets was most prominent in Tr1 cells, followed by Th1 cells, with minimal numbers of Tfh cells showing expression. The expression of NKG7 by Th1 and Tr1 cells is supported by previous studies[65–67, 96–98]. Additional experiments to address why NKG7 expression is higher in Tr1 cells than in Th1 cells are warranted. A potential explanation is that Tr1 cells represent more terminally differentiated Th1 cells[83], and this may result in the accumulation of NKG7 in this more differentiated cell subset. Alternatively, there may be specific signalling pathways involved in Tr1 cell differentiation that require NKG7. Hence, NKG7 may play an important role in balancing the anti-parasitic functions of Th1 cells with the tissue-protective roles of Tr1 cells, both of which are crucial for mounting an effective immune response during the blood stage malaria. The findings reported here suggest a dual role for NKG7 in both promoting anti-parasitic responses and dampening inflammation in order to protect tissue.

NKG7 has been recognised as a biomarker in various human diseases[74–76, 99–103], including different types of cancer[69–73, 104, 105]. Several human transcriptional studies have associated NKG7 with cytotoxic functions in CD4^+^ T cells[60–62, 95] and CD8^+^ [63, 64, 85–88]. Nevertheless, there has been a lack of comprehensive characterisation of NKG7 at the protein level in human diseases. Here, consistent with previous data, NKG7 was shown to be highly expressed by human cytotoxic and non-cytotoxic PBMCs from participants in CHMI caused by *P. falciparum* infection and uncomplicated malaria patients from disease endemic areas. NKG7 was expressed by populations of CD4^+^ and CD8^+^ cytotoxic T cells, as well as non-cytotoxic T cells. Notably, CD161 was associated with non-cytotoxic CD4^+^ and CD8^+^ T cells that also expressed high levels of NKG7, warranting further investigation. Additionally, *Plasmodium* species-specific differences in NKG7 expression were identified in patients with malaria. Further investigation is required to determine whether these NKG7-specific expression patterns are linked to specific outcomes, such as altered inflammation or immunity. Overall, these findings confirm that NKG7 is expressed by a range of CD4^+^ and CD8^+^ T cell populations in malaria, suggesting a variety of immunological roles following infection that are disease- and context-dependent.

There are several limitations with this study. Although experimental mouse models provide valuable insights, conclusions drawn from them cannot be directly extrapolated to the outcomes of malaria in humans. While certain similarities exist between rodent models or experimental human infection, and clinical infection, exact comparisons are not feasible, so care must be taken when making inferences. Furthermore, a limitation with the human samples examined was that they were restricted to cases of pre-symptomatic or uncomplicated malaria. It is also important to note that the CHMI datasets involved malaria-naïve volunteers, and outcomes in patients with malaria in disease endemic areas likely differ. Finally, all human datasets were constrained by samples size, limiting statistical power. Thus, careful consideration is required when drawing conclusions, and further investigation with larger samples sizes is warranted.

Overall, the findings in this study highlight the importance of NKG7 expressed by CD4^+^ and CD8^+^ T cells in both malaria disease and immunity, with the outcome influenced by a range of factors including the specific *Plasmodium* species responsible for the infection, the disease context and the organs involved. These data advance our understanding of host immune responses to malaria and suggest that NKG7 may hold promise as a potential therapeutic target to improve immune responses during malaria, as well as other parasitic diseases and inflammatory conditions.

## Materials and Methods

### Mice and human ethics

#### Mice

Experimental mice use was in accordance with the “Australian Code of Practice for the Care and Use of Animals for Scientific Purposes” (Australian National Health and Medical Research Council) and approved by the QIMR Berghofer Medical Research Institute Animal Ethics Committee (AEC; AEC reference number P2304; approval number: A1707-615M).

#### CHMI subjects

Human ethics approval was provided by the QIMR Berghofer Medical Research Institute Human Ethics Committee (HREC; HREC reference number P1479). Written informed consent was received from all participants[80].

#### Endemic country malaria patients

Ethics approval for the use of human samples was obtained from the QIMR Berghofer HREC (HREC reference number P3444), the Northern Territory Department of Health and Menzies School of Health Research ethics committee (HREC 2010-1431), and Medical Research and Ethics Committee, Ministry of Health, Malaysia (NMRR-10-754-6684 and NMRR-12-499-1203). Written informed consent was obtained from all adult study participant or, in the case of children, parents or legal guardians. The inclusion criteria included being 12 years of age or older.

### Experimental mice and human participant details

#### Mice

The experiments involved the use of female mice aged six week or older, unless otherwise specified. Mice were kept in group housing, with a maximum of six mice per cage. They were maintained in a pathogen-free environment at the QIMR Berghofer Medical Research Institute Animal Facility (Brisbane, Queensland, Australia). B6.*Cd45.1* (Ptprca) mice were obtained from the Animal Resources Centre (Perth, Western Australia, Australia). *C57BL/6J* (WT) mice were obtained from the Walter and Eliza Hall Institute (Melbourne, Victoria, Australia). All other mice were bred at QIMR, including B6J.*Nkg7*^−/−^, PbT-II, PbT-I and *Nkg7*-reporter (*Nkg7*-cre × mT/mG) lines. Transgenic PbT-II mice [78] were bred with B6J.*Nkg7*^−/−^ mice to produce *Nkg7*-deficient PbT-II mice (PbT-II^Δ*Nkg7*^; CD45.2^+^ CD45.1^−^) mice, and bred with B6.*Cd45.1* mice to produce PbT-II × B6.*Cd45.1* (PbT-II^WT^; CD45.1^+^ CD45.2^+^). Transgenic PbT-I mice [106] were bred with B6J.*Nkg7*^−/−^ mice to produce *Nkg7*-deficient PbT-I mice (PbT-I^Δ*Nkg7*^; CD45.1^−^ CD45.2^+^) mice, and bred with B6.*Cd45.1* mice to produce PbT-I × B6.*Cd45.1* (PbT-I^WT^; CD45.2^+^ CD45.1^+^).

#### CHMI subjects

PBMCs were obtained from volunteers who participated in a CHMI study involving *P. falciparum*. The primary investigation was a Phase 1b trial conducted at a single centre[80]. The trial aimed to evaluate the safety, tolerability, pharmacokinetic profile, and antimalarial activity of single doses of co-administered artefenomel and piperaquine phosphate against early *P. falciparum* blood-stage infection in healthy adult volunteers. Here, PBMCs from participants who consented for involvement in a study to analyse immunological responses were used. This included 8 individuals, 5/8 male, aged median 22, range 18-29.

#### Endemic country malaria patients

Blood was obtained from patients enrolled in a comparative clinical-pathophysiological study conducted at Queen Elizabeth Hospital (QEH) in Kota Kinabula, Sabah, Malaysia, spanning from September 2010 to October 2011 [107, 108]. This study incorporates analysis of PBMCs obtained from patients recruited in the primary study, including 5 individuals diagnosed with uncomplicated *P. falciparum* (2/5 male, median age 39, range 36-54) and 6 with uncomplicated *P. vivax* malaria (4/6 male, median age 42, range 16-54).

#### Plasmodium infection in mice

*Plasmodium* infections in mice were established using parasites that had undergone a single passage in C57BL/6J mice. 200 μl of transgenic *Pb*A (231c11) parasites, which were an in-house laboratory stock stored at −80 °C and expressing luciferase and GFP, were thawed at room temperature (RT). These parasites were then injected into a passage mouse via an intraperitoneal (i.p.) route. *Py*17XNL stabilates (in-house laboratory stock, frozen at −80 °C) were washed in NaCl and resuspended in 1 ml sterile PBS (1x) and 200 µl was injected via intravenous (i.v.) route into a passage mouse. Both the *Pb*A and *Py*17XNL passage mice were monitored with parasitemia checks from day 3 post infection (p.i.) and sacrificed when >1 % pRBCs. The passage mouse was euthanised and blood was obtained through cardiac puncture and diluted into RPMI/PS (containing 1 IU/ml heparin). The collected blood was then spun at 290 *g* for 7 min at RT. A hemocytometer (Pacific Laboratory Products) was used to enumerate RBCs. Parasite preparations were made, each containing 5×10^5^ pRBCs/ml. Subsequently, experimental mice were injected via i.v. route with 200 μl of this preparation, providing approximately 1×10^5^ pRBCs per mouse.

#### Quantification of Plasmodium parasitemia in mice

Passage and experimental mice were tail bled to collect two drops of blood into 250 μl RPMI/PS containing 1 IU/ml heparin. Next, 25 µl of this solution was incubated with SYTO 84 (Invitrogen; Life Technologies) and Hoechst 33342 (Sigma–Aldrich) in RPMI/PS for 22 min at RT. Then, each sample was quenched with 300 µl of RPMI/PS and acquired on a BD LSRFortessa through BD FlowJo version 10.0 (BD Biosciences).

#### Preparation of spleen single-cell suspensions

Mice were euthanized and a mid-sagittal incision was performed on the abdominal cavity and the spleen was carefully removed. The spleen’s weight was recorded, and it was then placed into 1 % fetal bovine serum (FBS) in PBS (1 % FBS/PBS). To isolate the cells, the spleens were mechanically mashed using the rear end of a 5 ml syringe plunger (Terumo Medical) through a 100 μm cell strainer (EASYstrainer; Greiner Bio-One). The resulting cell suspension was resuspended in 1 % FBS/PBS. Using an Eppendorf Centrifuge 5810R (Thermo Fisher Scientific), the cell suspension was then subsequently centrifuged at 350 *g*. To remove RBCs, the cell pellet was lysed by treating and agitating for 7 min at RT in Red Blood Cell Lysing Buffer Hybri-Max (Sigma–Aldrich). For enumeration, the cells were diluted in Trypan Blue Stain (Invitrogen) and 1x DPBS (Gibco). Further method details included in[53].

#### Preparation of brain mononuclear cells (MNCs)

Cardiac perfusions were carried out with DPBS before brains were removed. After extraction, the brain tissue was sectioned into smaller pieces and immersed in a solution containing 2 mg/ml collagenase in addition to 1 mg/ml deoxyribonuclease I (DNase I) obtained from bovine pancreas (both from Sigma–Aldrich). This mixture was prepared in 2 ml of Hanks’ Balanced Salt Solution (HBSS). Following this, the samples were placed on an Incu-Shaker Mini (Benchmark Scientific), set at 200 revolutions per minute (RPM) and maintained at 37 °C for a duration of 20 min. Following this incubation, the samples were homogenised utilising the back end of a 10 ml syringe plunger (Terumo Medical) through a 70 μm EASYstrainer cell strainer (Geiner Bio-One). 1 % FBS/PBS was used to rinse the single-cell suspensions from the brain. Using an Eppendorf Centrifuge 5810R (Thermo Fisher Scientific), the preparation was spun at 350 *g*. The supernatant was carefully removed, and 33 % Percoll Density Gradient Media (GE Healthcare) was used to resuspended the cell pellet. The next step involved spinning the preparation for 15 min at 575 *g* at RT, with the brake disengaged. Debris and any remaining supernatant were removed. The isolated MNCs were then incubated in 0.5 ml Red Blood Cell Lysis Buffer Hybri-Max (Sigma–Aldrich) for a period of 4 min at RT. After this incubation, MNCs were rinsed once again as detailed earlier and then cultured in 2x Monensin Solution (BioLegend) diluted in complete media. This incubation took place at 37 °C for a period of 3 h in the presence of 5 % CO_2_. For flow cytometry analysis of brain MNCs, staining panels incorporated the use of anti-mouse/human CD11b and anti-mouse F4/80 to exclude microglia from the analysis. Further method details included in[53].

#### Conventional and spectral flow cytometry

Flow cytometry staining protocols were carried out using Falcon 96-Well Clear Round Bottom Tissue Culture-Treated Cell Culture Microplates (Corning). For murine pre-stains, samples were incubated in 30 µl of complete media containing fluorescence-conjugated antibodies reactive against surface proteins for 30 min at 37 °C. 30 µl Zombie Aqua Fixable Viability Dye cocktail (BioLegend) or LIVE/DEAD Fixable Blue Dead Cell Stain Kit (Thermo Fisher) and TruStain FcX (BioLegend; anti-mouse CD16/32) were used for 15 min at RT to treat single-cell suspensions. During a 2 h stimulation at 37 °C, staining for CD107a (LAMP-1) was carried out with PMA/ionomycin or treatment with monensin (described later) by including 1 µl anti-Mouse CD107a (clone 1D4B from BioLegend) to 100 µl of the stimulation cocktail. Other PMA stimulations were carried out for a duration of 2.5 to 3 h. After treatment, cells were washed once with 1 % FBS/PBS and spun at 575 *g* for 1 min at 4 °C in an Eppendorf Centrifuge 5810R (Thermo Fisher Scientific). Subsequently, 30 µl of a master mix containing antibodies, conjugated with various fluorophores, reactive against surface proteins were used to stain samples for 30 min at 37 °C. After two washes with 1 % FBS/PBS staining buffer, samples were treated with 50 µl of fixation buffer for 20 min at RT. Fixation buffers used were from either the BD Cytofix Fixation Buffer Set (BD Biosciences) or the eBioscience Foxp3/Transcription Factor Staining Buffer Set (Thermo Fisher Scientific). Cells were then rinsed two times with wash buffers from the corresponding kits by spinning for 1 min at 953 *g* at RT. After this, cells were treated with 30 µl of a cocktail of antibodies, conjugated with various fluorophores, against intracellular proteins for 45 min at 37 °C. For human staining, 10^6^ PBMCs per participant, per time point, were aliquoted into plates. Samples were cultured with 50 µl of LIVE/DEAD Fixable Blue Dead Cell Stain Kit (diluted in 1x PBS; Thermo Fisher) at RT for 15 min. After this, cells were rinsed with 2 % FBS/PBS and spun for 5 min in an Eppendorf Centrifuge 5810R (Thermo Fisher Scientific). Subsequently, 25 µl of a first master mix of antibodies, conjugated with various fluorophores, reactive against surface proteins samples were used to stain samples for 15 min at RT. After this, 50 µl of a second master mix of antibodies, conjugated with various fluorophores, reactive against surface proteins was added to PBMCs for a further 30 min at RT. After two washes with 2 % FBS/PBS staining buffer, samples were treated with 100 µl of fixation buffer from the eBioscience Foxp3/Transcription Factor Staining Buffer Set (Thermo Fisher Scientific) for 20 min on ice. Cells were then washed twice with permeabilisation wash buffer. Following this, 50 µl of a master mix of antibodies, conjugated with various fluorophores, reactive against intracellular proteins cells were used to stain samples for 30 min on ice. All staining procedures were conducted in the dark. Prior to acquisition, samples were stored at 4 °C. Samples were acquired on a BD LSRFortessa (BD Biosciences) using BD FACSDiva version 8.0 or Cytek Aurora 5 laser using the SpectroFlo software package version 2.2 (Cytek Biosciences), then analysed on FlowJo version 10 OSX (FlowJo). GraphPad Prism 9 (version 9.4.0; GraphPad Software) was used for graphing and statistical analyses were conducted with statistical significance defined as P ≤ 0.05. Further method details included in[53].

Mouse flow cytometry markers used to define cell subsets can be found in Tables 1-4.

**Table 1.** Phenotypic flow cytometry markers for mouse datasets (Suppl Figure 1)

| Marker | Phenotype | Antibody<br>source | Catalogue<br>number | Marker | Phenotype | Antibody<br>source | Catalogue<br>number |
| --- | --- | --- | --- | --- | --- | --- | --- |
| TCRβ | T cell | BD<br>Biosciences | 612821 | TIGIT | Activation | BioLegend | 142106 |
| NK1.1 | NK cell | BioLegend | 156510 | CCR5 | Activation | BioLegend | 107016 |
| CD4 | T cell | BD<br>Biosciences | 563790 | CD107a | CTL | BioLegend | 121610 |
| CD8 | T cell | BioLegend | 100730 | GZMB | CTL | BioLegend | 515408 |
| CD45.1 | Congenetic<br>marker | BioLegend | 110706 | LAG-3 | Activation | BioLegend | 125219 |
| CD45.2 | Congenetic<br>marker | BD<br>Biosciences | 564616 | PD1 | Activation | BioLegend | 135240 |
| IFNγ | Cytokine | BD<br>Biosciences | 563854 | Perforin | CTL | BioLegend | 154406 |
| IL-10 | Cytokine | BioLegend | 505034 |  |  |  |  |

**Table 2.** Phenotypic flow cytometry markers for mouse datasets (Figure 2)

| Marker | Phenotype | Antibody<br>source | Catalogue<br>number | Marker | Phenotype | Antibody<br>source | Catalogue<br>number |
| --- | --- | --- | --- | --- | --- | --- | --- |
| TCRβ | T cell | BD<br>Biosciences | 612821 | PD1 | Activation | BioLegend | 135218 |
| CD8 | T cell | BioLegend | 100730 | TBET | Th1 cell | BioLegend | 644824 |
| CXCR5 | Tfh cell | BioLegend | 145506 | IFNγ | Cytokine | BD<br>Biosciences | 563854 |
| BCL6 | Tfh cell | BD<br>Biosciences | 563581 | IL-10 | Cytokine | BioLegend | 505034 |
|  |  |  |  | NK1.1 | NK cell | BioLegend | 156510 |

**Table 3.** Phenotypic flow cytometry markers for mouse datasets (Figure 3)

| Marker | Phenotype | Antibody<br>source | Catalogue<br>number | Marker | Phenotype | Antibody<br>source | Catalogue<br>number |
| --- | --- | --- | --- | --- | --- | --- | --- |
| TCRβ | T cell | BD Biosciences | 612821 | PD1 | Activation | BioLegend | 135218 |
| NK1.1 | NK cell | BioLegend | 156510 | eGFP | *Nkg7* expression |  |  |
| CD4 | T cell | BioLegend | 100548 | Td-Tomato | No *Nkg7* expression |  |  |
| CD8 | T cell | BioLegend | 100730 | IL-10 | Cytokine | BioLegend | 505034 |
| FOXP3 | Treg cell | BioLegend | 126408 | CXCR5 | Tfh cell | BioLegend | 145506 |
| CD45 | Lymphocytes | BD Biosciences | 564279 | CXCR6 | Th1 | BioLegend | 151119 |
| TBET | Th1 cell | BioLegend | 644824 | PD-1 | Activation | BioLegend | 135218 |
| IFNγ | Cytokine | BD Biosciences | 563854 | | | | |

**Table 4.** Phenotypic flow cytometry markers for mouse datasets (Figure 4)

| Marker | Phenotype | Antibody source | Catalogue number | Marker | Phenotype | Antibody source | Catalogue number |
| --- | --- | --- | --- | --- | --- | --- | --- |
| TCRβ | T cell | BD Biosciences | 612821 | PD1 | Activation | BioLegend | 135218 |
| NK1.1 | NK cell | BioLegend | 156510 | IFNγ | Cytokine | BD Biosciences | 563854 |
| CD4 | T cell | BioLegend | 100548 | IL-10 | Cytokine | BioLegend | 505034 |
| CD8 | T cell | BioLegend | 100730 | CXCR5 | Tfh cell | BioLegend | 145506 |
| FOXP3 | Treg | BioLegend | 126408 | CXCR6 | Th1 cell | BioLegend | 151104 |
| CD45.1 | Congenic marker | BioLegend | 110706 | TBET | Th1 cell | BioLegend | 644824 |
| CD45.2 | Congenic marker | BD Biosciences | 564616 |  |  |  |  |

Human flow cytometry markers used to define cell subsets can be found in Tables 5 & 6.

**Table 5.** Phenotypic flow cytometry markers for human datasets (Figure 5 and 6)

| Marker | Phenotype | Antibody source | Catalogue number | Marker | Phenotype | Antibody source | Catalogue number |
| --- | --- | --- | --- | --- | --- | --- | --- |
| CD16 | Monocyte | BD Biosciences | 612944 | CXCR3 | Th1 cell | BD Biosciences | 565223 |
| CD56 | NK cell | BioLegend | 318348 | CD8 | T cell | BioLegend | 344734 |
| GZMB | CTL | BioLegend | 515408 | HLA-DR | Activation | BD Biosciences | 335796 |
| CD107a | CTL | BD Biosciences | 562622 | CD14 | Monocyte | BD Biosciences | 612902 |
| KLRG1 | CTL | Thermo Fisher | 46-9488-42 | CD49b | Tr1 cell | Thermo Fisher | 11-0498-42 |
| CD4 | T cell | BD Biosciences | 612913 | CD38 | Activation | BD Biosciences | 563964 |
| CD25 | Treg cell | BioLegend | 356146 | CD45RA | TEMRA, Naive | BioLegend | 304134 |
| CCR4 | Th2 cell | BioLegend | 359416 | CCR7 | TCM, Naive | BD Biosciences | 561144 |
| PD1 | Activation | BD Biosciences | 561272 | CCR6 | Th17 cell | BD Biosciences | 563922 |
| CD3 | T cell | Thermo Fisher | 58-0032-82 | CXCR5 | Tfh cell | BioLegend | 356942 |
| CD127 | Treg cell | BD Biosciences | 565185 | CD19 | B cell | BioLegend | 302274 |
| NKG7 | NKG7 | Beckman-Coulter | IM3293 |  |  |  |  |

**Table 6.** Cytotoxic flow cytometry markers for endemic dataset (Figure 6 and Supplementary Figure 4)

| Marker | Phenotype | Antibody source | Catalogue number | Marker | Phenotype | Antibody source | Catalogue number |
| --- | --- | --- | --- | --- | --- | --- | --- |
| CD3 | T cell | BD Biosciences | 612893 | CD4 | T cell | BD Biosciences | 612912 |
| GNLY | CTL | BD Biosciences | 558254 | CD127 | MPEC | BioLegend | 351308 |
| GZMB | CTL | BioLegend | 372204 | HLA-DR | Activation | BioLegend | 307672 |
| NKG7 | NKG7 | Beckman-Coulter | IM3293 | CD16 | NK cell | BD Biosciences | 563785 |
| CD45RA | TEMRA, Naive | BioLegend | 304138 | TCR$\gamma\delta$ | T cell | BD Biosciences | 564157 |
| CD27 | TCM, Naive | BioLegend | 356418 | PRF1 | CTL | BD Biosciences | 563763 |
| CD8 | T cell | BioLegend | 344740 | TCRVδ2 | T cell | BioLegend | 331420 |
| CD56 | NK cell | BD Biosciences | 612766 | KLRG1 | SLEC | Thermo Fisher | 25-9488-42 |
| CD25 | Activation | BD Biosciences | 565106 | CD161 | Activation | BD Biosciences | 563864 |
| CD57 | Differentiation | BioLegend | 359608 |  |  |  |  |

**Table 7.** Resources and equipment[109].

| REAGENT or RESOURCE | SOURCE | IDENTIFIER |
| --- | --- | --- |
| Human TruStain FcX | BioLegend | Cat#422302 |
| TruStain FcX (α-mouse CD16/32) Antibody (clone: 93) | BioLegend | Cat#101320 |
| TruStain FcX PLUS α-mouse CD16/32 (clone: S17011E) | BioLegend | Cat#156604 |
| **Chemicals** |  |  |
| 2-Mercaptoethanol | Sigma-Aldrich, St. Louis MI, USA | Cat#M6250 |
| BD GolgiStop Protein Transport Inhibitor (Containing Monensin) | BD Biosciences | Cat#554724 |
| Benzonase Nuclease |  | Cat#9025-65-4 |
| Brilliant Stain Buffer Plus | BD Biosciences | Cat#566385 |
| Dulbecco's Phosphate Buffered Saline (DPBS) | Gibco, Life Technologies, Carlsbad CA, USA | Cat#14190250 |
| Ficoll-Paque PLUS | GE Healthcare, Little Chalfont, Buckinghamshire, U.K. | Cat#17-1440-03 |
| Hoeschst 33342 (bisbenzimidazole H 333342 trihydrochloride Cas#23491-52-3) | Sigma-Aldrich | Cat#B2261 |
| Ionomycin calcium salt from *Streptomyces conglobatus* | Sigma-Aldrich | Cat#I0634-1MG |
| MEM Non-Essential Amino Acids Solution (100X) | Gibco, Life Technologies | Cat#11140050 |
| Penicillin-Streptomycin (10,000 U/ml) | Gibco, Life Technologies | Cat#14140122 |
| Phorbol 12-myristate 13-acetate (PMA) | Sigma-Aldrich | Cat#P8139-1MG |
| Red Blood Cell Lysing Buffer Hybri-Max | Sigma-Aldrich | Cat#R7757 |
| Syto 84 Orange Fluorescent Nucleic Acid Stain | Invitrogen, Carlsbad<br>CA, USA | Cat#S11365 |
| Trypan Blue Stain (0.4%) for use with Countess<br>Automated Cell Counter | Invitrogen | Cat#T10282 |
| UltraPure DNase/RNase-Free Distilled Water | Invitrogen | Cat#10977015 |
| **Commercial Kits** |  |  |
| 7-AAD Cell Viability |  |  |
| Annexin V PE Apoptosis Detection Kit I | BD Biosciences | Cat#559763 |
| CD4<sup>+</sup> T Cell Isolation Kit, mouse | Miltenyi Biotec | Cat#130-104-454 |
| CD8<sup>+</sup> T Cell Isolation Kit, mouse | Miltenyi Biotec | Cat#130-104-075 |
| EasySep Mouse CD4<sup>+</sup> Cell Isolation Kit | STEMCELL | Cat#19852 |
| EasySep Mouse CD8<sup>+</sup> Cell Isolation Kit | STEMCELL | Cat#19853 |
| eBioscience Foxp3/Transcription Factor Staining<br>Buffer Set | eBioscience, Thermo<br>Fisher Scientific | Cat#00-5523-00 |
| LIVE/DEAD Fixable Blue Dead Cell Stain Kit | Thermo Fisher | Cat#L34962 |
| NEBNext Single Cell/Low Input RNA Library Prep<br>Kit for Illumina | New England BioLabs | Cat#E6420S |
| RNase-Free DNase Set | QIAGEN | Cat#79254 |
| RNeasy Micro Kit | QIAGEN | Cat#74004 |
| RNeasy Mini Kit | QIAGEN | Cat#74104 |
| SYTOX Blue Dead Cell Stain | Thermo Fisher | Cat#S34857 |
| Zombie Aqua Fixable Viability Kit | BioLegend | Cat#423102 |
| Zombie NIR Fixable Viability Kit | BioLegend | Cat#423106 |
| ZymoScript RT PreMix | Zyma Research | Cat#R3012 |
| **Disposables** |  |  |
| 10cc/ml Syringe | Terumo Medical | Cat#SS-10S |
| 1cc/ml Tuberculin Syringe | Terumo Medical,<br>Somerset NJ, USA | Cat#SS-01T |
| 26G x ½" Needle | Terumo Medical | Cat#NN-2613R |
| 5cc/ml Syringe | Terumo Medical | Cat#SS-05S |
| BD Vacutainer Lithium Heparin (LH) 170 I.U. Plus Blood Collection Tubes | BD Biosciences | Cat#367526 |
| Corning cells strainer, size 100um | Sigma-Aldrich | Cat#CLS43175 |
| Countess Cell Counting Chamber Slides | Invitrogen | Cat#C10228 |
| EASYstrainer 100um | Greiner Bio-One,<br>Kremsmunster,<br>Austria | Cat#542000 |
| Falcon 96-well Clear Round Bottom Tissue Culture (TC)-Treated Cell Culture Microplates | Corning Inc., Corning<br>NY, USA | Cat#353077 |
| **Equipment** |  |  |
| Bio-Rad CFX384 Touch Real-Time PCR Detection System | Bio-Rad |  |
| Countess II FL Automated Cell Counter | Invitrogen | Cat#AMQAP1000 |
| Cytek Aurora | Cytek | Special order<br>research product |
| Eppendorf Centrifuge | Fisher Scientific<br>Thermo Fisher<br>Scientific | Cat#05-413 |
| FACSARIA II | BD Biosciences | Special order<br>research product |
| LSRFortessa: 4 laser configuration | BD Biosciences | Special order<br>research product |
| LSRFortessa: 5 laser configuration | BD Biosciences | Special order<br>research product |
| Olympus CX31 Biological Microscope | Olympus Life Science,<br>Shinjuku, Tokyo,<br>Japan | Model#CX31 |
| **Experimental Models** |  |  |
| Mouse: B6.129(Cg)-*Gt(ROSA)26Sor<sup>tm4</sup>(ACTB-tdTomato,-EGFP)*<sup>Luo</sup>/J (B6.mTmG) | Liqun Luo, Howard<br>Hughes Medical<br>Institute, Stanford | RRID:<br>IMSR_JAX:007676 |
|  | University, Stanford<br>CA, USA |  |
| Mouse: B6.SJL-Ptprc<sup>a</sup> Pepc<sup>b</sup>/BoyJ (B6.Cd45.1) | Animal Resources<br>Centre, Canning Vale,<br>WA, Australia | RRID:<br>IMSR_JAX:002014 |
| Mouse: B6.Tg(*Nkg7-cre*)/J (B6J.*Nkg7-cre*) | Melbourne Advanced<br>Genome Editing<br>Centre (MAGEC), The<br>Walter and Eliza Hall<br>Institute (WEHI),<br>Parkville VIC, Australia | Commissioned |
| Mouse: B6J.*Nkg7* ko | QIMRB, Brisbane,<br>Australia | RRID:<br>IMSR_KOMP:VG114<br>45-1.1-Vlcg |
| Mouse: *C57BL.6J* | WEHI, Kew VIC,<br>Australia | RRID:<br>IMSR_JAX:000664 |
| Mouse: Transgenic PbT-I | QIMRB, Brisbane,<br>Australia | Reference[106] |
| Mouse: Transgenic PbT-II | QIMRB, Brisbane,<br>Australia | Reference[78] |
| *Plasmodium berghei*: PbA | QIMRB, Brisbane,<br>Australia | Reference[110] |
| *Plasmodium yoelii*: Py17XNL | QIMRB, Brisbane,<br>Australia |  |
| **Media** |  |  |
| Dulbecco's Modified Eagle Medium (DMEM),<br>high glucose, pyruvate | Gibco, Life<br>Technologies | Cat#11995073 |
| Fetal Bovine Serum, certified, US origin | Gibco, Life<br>Technologies | Cat#16000044 |
| Roswell Park Memorial Institute Medium (RPMI)<br>1640 Medium | Gibco, Life<br>Technologies | Cat#11875119 |
| **Oligonucleotides/Recombinant DNA** |  |  |
| Hs00366585_g1 human NKG7 FAM-MGB TaqMan probe | Thermo Fisher | Cat#4453320 |
| Hs03003631_g1 Human 18S VIC-MGB TaqMan probe | Thermo Fisher | Cat#4453320 |
| **Software and Packages** |  |  |
| EdgeR | https://bioconductor.org/packages/release/bioc/html/edgeR.html |  |
| FlowJo, v10 OSX | FlowJo, LLC.<br>https://www.flowjo.com |  |
| Ggplot2 | https://www.gsea-msigdb.org/gsea/index.jsp |  |
| GraphPad Prism 9, v9.4.0 | GraphPad Software<br>https://www.graphpad.com |  |
| Morpheus | https://software.broadinstitute.org/morpheus/ |  |
| OMIQ | OMIQ, by Dotmatics<br>https://www.omiq.ai |  |
| RStudio Desktop | https://posit.co |  |
| The R Project for Statistical Computing | https://www.r-project.org | RRID: SCR_001905 |

#### PMA/ionomycin restimulation

Cells were cultured in complete media supplemented with 25 ng/ml PMA (Sigma–Aldrich) and 1 μg/ml (1.33 nM) ionomycin calcium salt (Sigma–Aldrich), with the addition of 2x monensin solution (BioLegend). The PMA/ionomycin restimulation process was carried out over a 2-3 h duration at 37 °C in the presence of 5 % CO_2_. Further method details included in[53].

#### Isolation of PbT-II and PbT-I cells by FACS

PbT-II cells were enriched prior to FACS by using either the CD4+ T cell Isolation Kit, mouse (Miltenyi Biotec) or EasySep Mouse CD4+ T Cell Isolation Kit (STEMCELL) following the manufacturer’s guidelines. PbT-I cells were enriched prior to FACS by using either the CD8+ T cell Isolation Kit, mouse (Miltenyi Biotec) or EasySep Mouse CD8+ T Cell Isolation Kit (STEMCELL) following the manufacturer’s guidelines. PbT-II and PbT-I cells were stained for surface proteins prior to isolation by cell sorting for RNA-sequencing and real-time quantitative PCR (RT-qPCR). The surface staining included use of anti-mouse CD90.2 (BV605; 53-2.1), anti-mouse CD45.2 (PE; 104), anti-mouse CD45.1 (FITC; A20) and either anti-mouse CD4 (APC; GK1.5) for PbT-II cells or anti-mouse CD8 (APC; 53-6.7) for PbT-I cells (all from BioLegend). Dead cells were omitted during flow cytometry with a positive stain for SYTOX Blue Dead Cell Stain. Cells were sorted on a BD FACSAria II or BD FACSAria III (Becton Dickinson), treated in buffer RLT (Qiagen) and stored at −80 °C. Furhter method details included in[53].

#### Adoptive cell transfer of PbT-II and PbT-I cells

PbT-II cells were isolated from the spleens of PbT-II^WT^ (CD45.1 x PbT-II) and PbT-II^Δ*Nkg7*^ mice using either the CD4^+^ T cell Isolation Kit, mouse (Miltenyi Biotec) or EasySep Mouse CD4+ T Cell Isolation Kit (STEMCELL) following the manufacturer’s guidelines. PbT-I cells were isolated from the spleens of PbT-I^WT^ (CD45.1 x PbT-I) and PbT-I^Δ*Nkg7*^ mice using either the CD8a^+^ T Cell Isolation Kit, mouse (Miltenyi Biotec) or EasySep Mouse CD8^+^ T Cell Isolation Kit (STEMCELL) following the manufacturer’s guidelines. PbT-II^WT^ and PbT-II^Δ*Nkg7*^ cells were enumerated and pooled in equal numbers to a final concentration of 1×10^6^ cells/ml. Then, 1 day before infection with *Pb*A or *Py*17XNL, 200 μl (2×10^5^ cells) was injected via i.v. route into the tail vein of B6.*Cd45.1* recipient mice. Similarly, PbT-I^WT^ and PbT-I^Δ*Nkg7*^ cells were enumerated and pooled in equivalent numbers to a final concentration of 5×10^6^ cells/ml. Then, 200 μl (1×10^6^ cells) were injected via i.v. route into the tail vein of B6.*Cd45.1* recipient mice 1 day before infection with *Pb*A. Additional method details included in[53].

#### HEK293T transient transfection

HEK293T cells were plated at 0.5×10^6^ in 60mm plate (Falcon) with 2 ml DMEM (Gibco) supplemented with 10% heat inactivated FCS (Gibco) for 48 hours at 37°C 5% C0_2_ or until 50-60% confluent. NKG7-GFP reporter cells were generated by adding a transfection mix of RD114 envelope plasmid (1.5 μg), PeqPam Gag-pol plasmid (2 μg) and vector plasmid eGFP-T2A-NKG7 (6.25 μg) with 19 μl FuGENE 6 transfection reagent (Promega) in a final volume of 470 μl of DMEM, dropwise to the cells with gentle mixing. After transfection cells were incubated for 72 hours before flow cytometry analysis.

#### NK92 Cell culture

NK92 human NK cell line (Malignant non-Hodgkin’s lymphoma) were cultured in RPM1640 (Gibco) supplemented with 10% heat inactivated FCS (Gibco), 1x Glutamax (Gibco), 100 units/ml Penicillin-Streptomycin (Gibco) and 1000 units/ml recombinant human IL-2 (Miltenyi). Cells were incubated at 37°C 5% C0_2_ and were passaged every 48 hours in 10 ml volume at a density of 2-5×10^5^ cells/ml.

#### Nucleofection and CRISPR Cas9

Gene editing was performed on 5×10^5^ NK92 cells as previously described (Huang RS). Briefly, for CAS9-RNP preparation 2ul of CAS9-NLS (40 μM) (QB3 Macrolab) were mixed with 2.4ul of human *NKG7* sgRNA sequence: CATGGCCACCACCATGGAGA (100 μM) (Synthego) and incubated for 15 minutes at 37°C. NK92 cells were washed once in dPBS (Gibco) and suspended in 20ul of freshly prepared nucleofection buffer containing 150 mM sodium phosphate buffer (pH 7.2), 5 mM KCl, 15 mM MgCl_2_, 15 mM HEPES and 50 mM mannitol (SIGMA), cells were added to the CAS9-RNP construct and 24.4 μl volume was transferred to a 16 well nucleofection strip (Lonza) for pulsing with program CM-137 on the Amaxa 96 well shuttle (Lonza). Nucleofected cells were transferred to a 24 well plate (Falcon). One ml of warm culture media was added and cells were incubated for 96 hours at 37°C 5% C0_2_, followed by flow cytometry staining.

#### RNA extraction

NK92 cells were preserved in RLT buffer and subsequently homogenised in QIAshredder columns (both from Qiagen). The RNA extraction process was performed using either the RNeasy Mini Kit or RNeasy Micro Kit (both from Qiagen), following the manufacturer’s guidelines. RNA integrity was assessed using the RNA 6000 Pico Kit (Agilent Technologies) (Performed at QIMR Berghofer sequencing facility). Subsequently, the isolated RNA for RT-qPCR was reverse transcribed into cDNA using the iScript cDNA Synthesis Kit (Bio-Rad) following the manufacturer’s instructions.

#### RT-qPCR

Taqman Gene Expression Assays (Life Technologies) were utilised in conjunction with GoTaq Probe qPCR Master Mix (Promega Corporation) following the recommended cycling settings suggested by the manufacturer. Reactions were carried out in a volume of 10 μl, including 1 μl of template cDNA. RT-qPCR was conducted in HardShell 384-Well Plates (Bio-Rad). Microseal ‘B’ PCR Plate Sealing Film (Bio-Rad) was used to seal the plates. CFX96 Touch Real-Time PCR Detection System (Bio-Rad) was used to run the TaqMan Gene Expression Assays. To determine relative gene expression levels, the comparative CT method was employed, using mean CT values of *NKG7*, compared to *18S* (for TaqMan Gene Expression Assays).

### Statistical analysis

Statistical analysis was performed using GraphPad Prism 6 (GraphPad Software, Boston, MA). Analysis of parasite area under the curve (AUC) in mouse experiments was performed using a Mann-Whitney test, while differences in mouse flow cytometry data was tested using a two-way ANOVA with Sidak’s multiple comparisons test or one-way ANOVA with Tukey’s multiple comparisons test, as appropriate. Differences in PbTII cell frequencies in mouse experiments were assessed using a Wilcoxon test. Analysis of human cell clusters was performed using Edge R, while differences in immune molecule expression by clusters was assessed by one-way ANOVA with Tukey’s multiple comparisons test.

## Data availability statement

All data are available in the main text and supplemental materials, and values for all data points in graphs, are shown in Table 8.

## Acknowledgements

QIMR-Berghofer, Burnet Institute and the University of Melbourne acknowledge the traditional custodians of the lands where they are located, the Turrbal and Jagera people (Brisbane), and the Boonwurrung and Wurundjeri Woi-wurrong people (Melbourne).

We thank the participants involved in the CHMI studies and all study clinicians and support staff at QPharm (Brisbane, Australia). These studies were funded by the Medicines for Malaria Venture.

This work was supported by the National Health and Medical Research Council of Australia (program grant 1132975 C.R.E.); Senior Research Fellowships C.R.E. (1154265), Career Development Award 1141278, Project Grant 1125656, and Ideas Grant 1181932 to MJB); the CSL Centenary Fellowship and the Snow Medical Foundation Fellowship 2022/SF167 to M.J.B. The Burnet Institute is supported by the NHMRC for Independent Research Institutes Infrastructure Support Scheme and the Victorian State Government Operational Infrastructure Support.

## Supplementary Figures

**Supplementary Figure 1.**
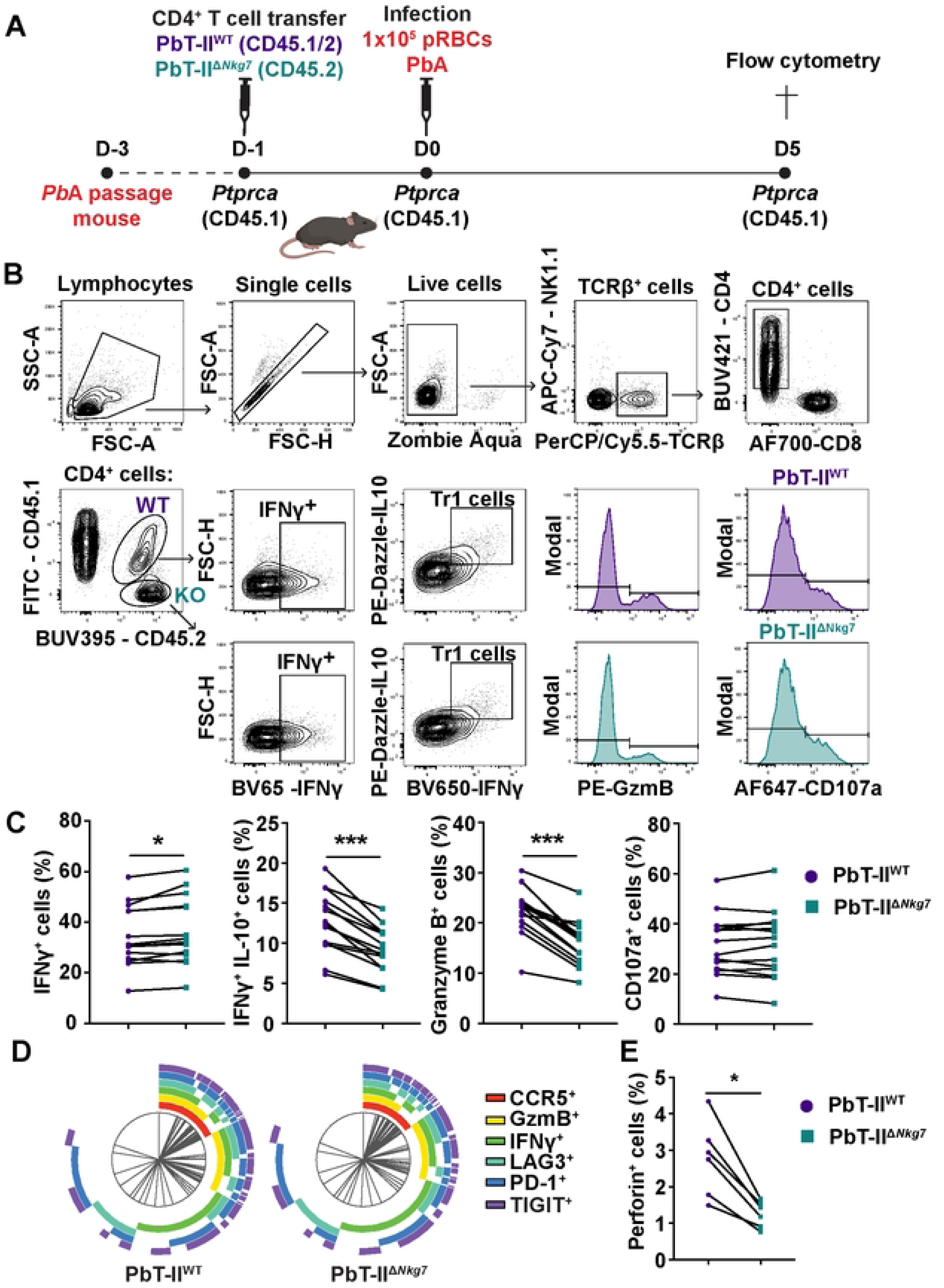
NKG7 has a cell-intrinsic role in the development of CD4^+^ T cell responses at day five of *Plasmodium berghei* ANKA (*Pb*A) infection. Congenic *Ptprca* (CD45.1) mice co-transferred with PbT-II^WT^ and PbT-II^ΔNkg7^ cells were infected with *Pb*A, causing experimental cerebral malaria (ECM). Schematic showing the experimental timeline (A). The gating strategy used for the experiment (B). The four graphs indicate frequencies of IFNγ^+^, Tr1 (IFNγ^+^ IL-10^+^), Granzyme B^+^ and CD107a^+^ PbT-II^WT^ and PbT-II^ΔNkg7^ cells at the onset of ECM (day 5) after stimulation with phorbal 12-myristate 13-acetate (PMA) and ionomycin *ex vivo*, in the presence of monensin (C). Data is pooled from two independent experiments. A Wilcoxon test were used to determine statistical significance, where * P < 0.05, *** P < 0.001. n = 14 infected mice. The SPICE graphs depict differences in co-expression of cellular markers between co-transferred transgenic PbT-II^WT^ and PbT-II^ΔNkg7^ cells in the spleen of infected mice (at ECM onset (day 5)) after treatment with phorbal 12-myristate 13-acetate (PMA) and ionomycin, in the presence of monensin (D). Data is from one experiment. n = 6 infected mice. The graph depicts the frequencies of Perforin^+^ PbT-II^WT^ and PbT-II^ΔNkg7^ cells in the spleen of infected mice (at peak ECM being day 5 p.i.) after treatment with phorbal 12-myristate 13-acetate (PMA) and ionomycin, in the presence of monensin (E). A Wilcoxon test was utilised to calculate statistical significance, where * P < 0.05. Data represents one experiment. n = 6 infected mice.

**Supplementary Figure 2.**
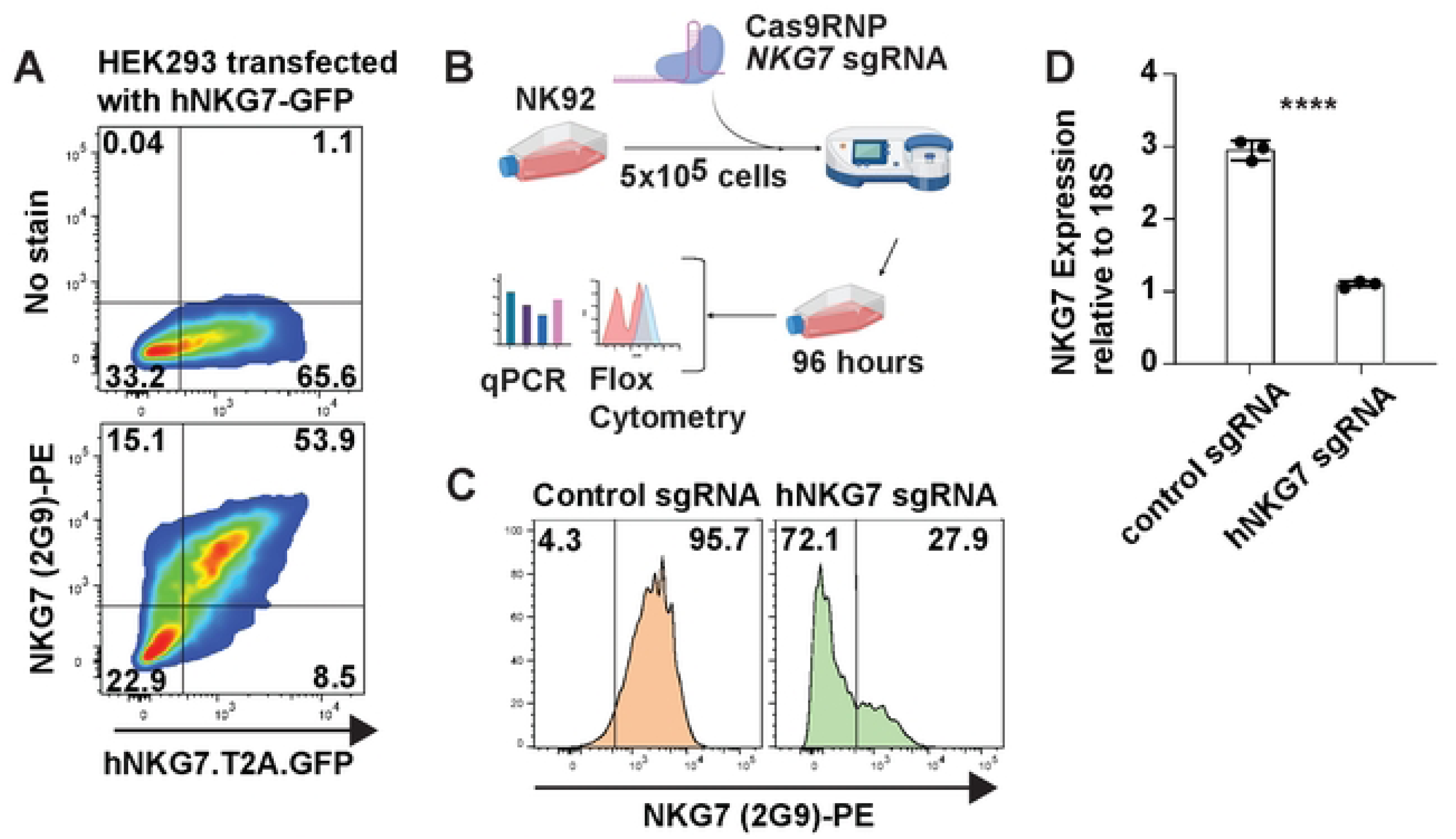
The 2G9 monoclonal antibody recognises human NKG7. HEK293 cells were transiently transfected with human NKG7 fused to green fluorescent protein. Flow cytometry analysis of cells showing GFP expression and 2G9 binding (lower panel), compared with no antibody stain (upper panel). Percentage of cells are indicated in each quadrant (A). The work flow for *NKG7* gene editing with single guide (sg)RNA and CRISPR/Cas9 in the human NK92 cell line (B). 2G9 binding was assessed by flow cytometry in control NK92 cells (left panel) and *NKG7* edited cells (right panel) with percentages of positive and negative cells indicated (C). NKG7 gene expression, relative to 18S RNA, was assessed by qPCR in control NK92 cells and *NKG7* edited cells, as indicated (D). Significance was assessed using an unpaired t-test, where **** P < 0.001.

**Supplementary Figure 3.**
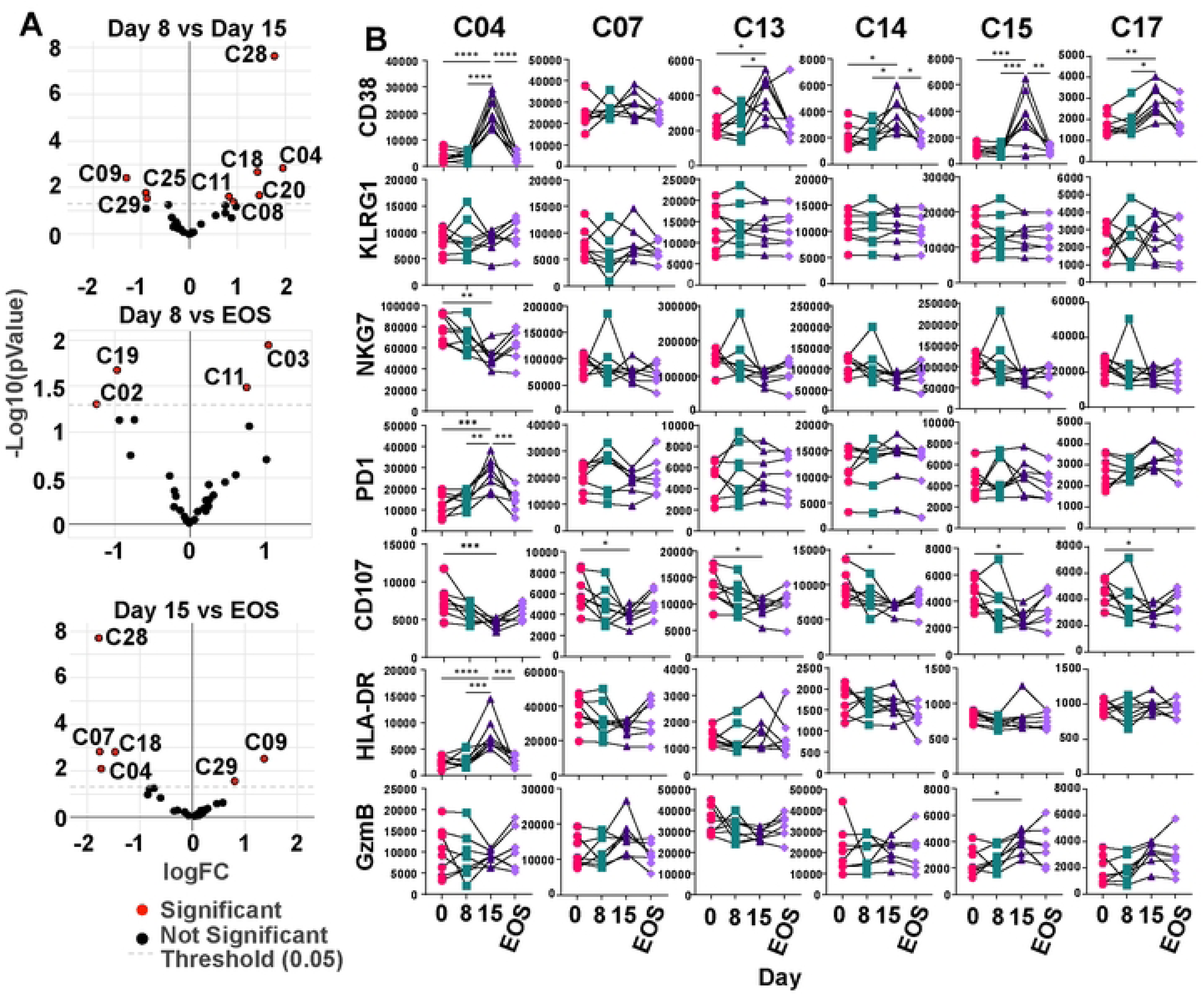
NKG7-positive CD4^+^ T cells had an activated phenotype at day 15 post *Plasmodium falciparum* infection. Peripheral blood mononuclear cells were acquired by spectral flow cytometry and analysed with dimensionality reduction using unsupervised FLOWSOM clustering in OMIQ software. PBMCs were divided into 30 clusters and clusters that were significantly different in abundance between day 8 and 15, day 8 and end of study (EOS), and day 15 and EOS are indicated with red circles. Day 0 (n = 8), day 8 (n = 8), day 15 (n= 8), EOS (n = 7). Significance assessed using edgeR (A). The expression of CD38, Killer cell lectin-like receptor G1 (KLRG1), NKG7, programmed cell death protein 1 (PD1), CD107a, HLA-DR and Granzyme B (GZMB) was measured, and the mean fluorescence intensity (MFI) of cellular markers is shown (B). Day 0 (n = 8), day 8 (n = 8), day 15 (n= 8), end of study (EOS; n = 7). ** P < 0.01, *** P < 0.001, **** P < 0.0001; significance assessed by one-way ANOVA with Tukey’s multiple comparisons test.

**Supplementary Figure 4.**
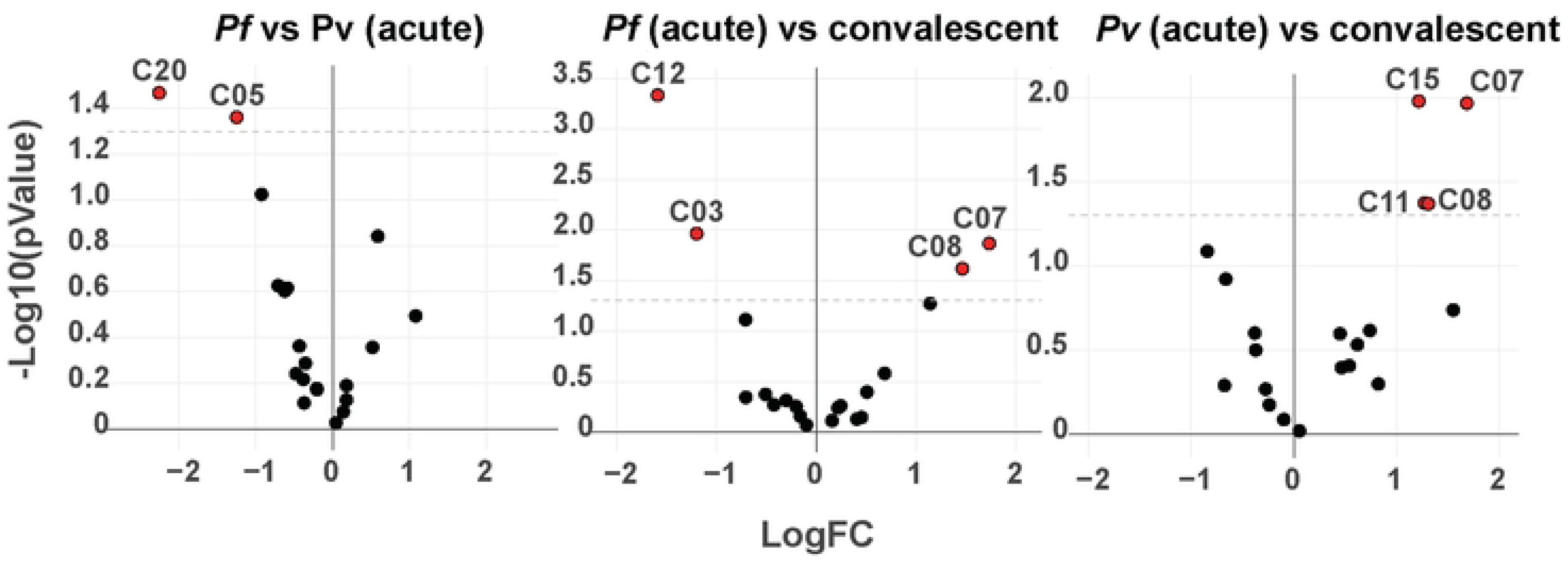
NKG7-expresing CD4^+^ T cell clusters were significantly more abundant during acute uncomplicated *Plasmodium vivax* (*Pv*) infection when compared to acute *P. falciparum* (*Pf*) infection. Peripheral blood mononuclear cells were acquired by spectral flow cytometry and analysed with dimensionality reduction using unsupervised FLOWSOM clustering in OMIQ software. PBMCs were divided into 20 clusters and clusters that were significantly different in abundance between Acute *Pf* and Acute *Pv*, Acute *Pf* vs Convalescence and Acute *Pv* vs Convalescence are indicated with red circles. Acute *Pf* (n = 5), acute *Pv* (n = 5), convalescence (n= 8). Significance assessed using edgeR.

## References

1. WHO. World malaria report 2024. Geneva: World Health Organization, 2024.

2. Organization GWH. World malaria report 2022. 2022 Contract No.: Licence: CC BY-NC-SA 3.0 IGO.

3. Boyle MJ, Engwerda CR, Jagannathan P. The impact of Plasmodium-driven immunoregulatory networks on immunity to malaria. Nat Rev Immunol. 2024. Epub 2024/06/12. doi: 10.1038/s41577-024-01041-5. PubMed PMID: 38862638.

4. Carter R. Studies on enzyme variation in the murine malaria parasites Plasmodium berghei, P. yoelii, P. vinckei and P. chabaudi by starch gel electrophoresis. Parasitology. 1978;76(3):241–67. Epub 1978/06/01. doi: 10.1017/s0031182000048137. PubMed PMID: 351525.

5. Wykes MN, Good MF. What have we learnt from mouse models for the study of malaria? Eur J Immunol. 2009;39(8):2004–7. Epub 2009/08/13. doi: 10.1002/eji.200939552. PubMed PMID: 19672886.

6. Taylor-Robinson AW. Regulation of immunity to Plasmodium: implications from mouse models for blood stage malaria vaccine design. Exp Parasitol. 2010;126(3):406–14. Epub 2010/02/09. doi: 10.1016/j.exppara.2010.01.028. PubMed PMID: 20138874.

7. Wassmer SC, de Koning-Ward TF, Grau GER, Pai S. Unravelling mysteries at the perivascular space: a new rationale for cerebral malaria pathogenesis. Trends Parasitol. 2024;40(1):28–44. Epub 2023/12/09. doi: 10.1016/j.pt.2023.11.005. PubMed PMID: 38065791; PubMed Central PMCID: PMCPMC11072469.

8. Ishizuka AS, Lyke KE, DeZure A, Berry AA, Richie TL, Mendoza FH, et al. Protection against malaria at 1 year and immune correlates following PfSPZ vaccination. Nat Med. 2016;22(6):614–23. Epub 2016/05/10. doi: 10.1038/nm.4110. PubMed PMID: 27158907.

9. Roestenberg M, McCall M, Hopman J, Wiersma J, Luty AJ, van Gemert GJ, et al. Protection against a malaria challenge by sporozoite inoculation. N Engl J Med. 2009;361(5):468–77. Epub 2009/07/31. doi: 10.1056/NEJMoa0805832. PubMed PMID: 19641203.

10. Mordmuller B, Surat G, Lagler H, Chakravarty S, Ishizuka AS, Lalremruata A, et al. Sterile protection against human malaria by chemoattenuated PfSPZ vaccine. Nature. 2017;542(7642):445-9. Epub 2017/02/16. doi: 10.1038/nature21060. PubMed PMID: 28199305.

11. Seder RA, Chang LJ, Enama ME, Zephir KL, Sarwar UN, Gordon IJ, et al. Protection against malaria by intravenous immunization with a nonreplicating sporozoite vaccine. Science. 2013;341(6152):1359-65. Epub 2013/08/10. doi: 10.1126/science.1241800. PubMed PMID: 23929949.

12. Kumar R, Loughland JR, Ng SS, Boyle MJ, Engwerda CR. The regulation of CD4(+) T cells during malaria. Immunol Rev. 2020;293(1):70–87. Epub 2019/11/02. doi: 10.1111/imr.12804. PubMed PMID: 31674682.

13. Troye-Blomberg M, Andersson G, Stoczkowska M, Shabo R, Romero P, Patarroyo ME, et al. Production of IL 2 and IFN-gamma by T cells from malaria patients in response to Plasmodium falciparum or erythrocyte antigens in vitro. J Immunol. 1985;135(5):3498–504. Epub 1985/11/01. PubMed PMID: 3930605.

14. Su Z, Stevenson MM. Central role of endogenous gamma interferon in protective immunity against blood-stage Plasmodium chabaudi AS infection. Infect Immun. 2000;68(8):4399–406. Epub 2000/07/19. doi: 10.1128/iai.68.8.4399-4406.2000. PubMed PMID: 10899836; PubMed Central PMCID: PMCPMC98333.

15. Blanchette J, Jaramillo M, Olivier M. Signalling events involved in interferon-gamma-inducible macrophage nitric oxide generation. Immunology. 2003;108(4):513–22. Epub 2003/04/02. doi: 10.1046/j.1365-2567.2003.01620.x. PubMed PMID: 12667213; PubMed Central PMCID: PMCPMC1782926.

16. Oakley MS, Sahu BR, Lotspeich-Cole L, Majam V, Thao Pham P, Sengupta Banerjee A, et al. T-bet modulates the antibody response and immune protection during murine malaria. Eur J Immunol. 2014;44(9):2680–91. Epub 2014/07/23. doi: 10.1002/eji.201344437. PubMed PMID: 25047384.

17. McCall MB, Hopman J, Daou M, Maiga B, Dara V, Ploemen I, et al. Early interferon-gamma response against Plasmodium falciparum correlates with interethnic differences in susceptibility to parasitemia between sympatric Fulani and Dogon in Mali. J Infect Dis. 2010;201(1):142–52. Epub 2009/11/26. doi: 10.1086/648596. PubMed PMID: 19929378.

18. Tubo NJ, Jenkins MK. CD4+ T Cells: guardians of the phagosome. Clinical microbiology reviews. 2014;27(2):200–13. doi: 10.1128/CMR.00097-13. PubMed PMID: 24696433; PubMed Central PMCID: PMC3993097.

19. Portugal S, Tipton CM, Sohn H, Kone Y, Wang J, Li S, et al. Malaria-associated atypical memory B cells exhibit markedly reduced B cell receptor signaling and effector function. Elife. 2015;4. Epub 2015/05/09. doi: 10.7554/eLife.07218. PubMed PMID: 25955968; PubMed Central PMCID: PMCPMC4444601.

20. Rivera-Correa J, Mackroth MS, Jacobs T, Schulze Zur Wiesch J, Rolling T, Rodriguez A. Atypical memory B-cells are associated with Plasmodium falciparum anemia through anti-phosphatidylserine antibodies. Elife. 2019;8. Epub 2019/11/13. doi: 10.7554/eLife.48309. PubMed PMID: 31713516; PubMed Central PMCID: PMCPMC6853636.

21. Boyle MJ, Jagannathan P, Bowen K, McIntyre TI, Vance HM, Farrington LA, et al. The Development of Plasmodium falciparum-Specific IL10 CD4 T Cells and Protection from Malaria in Children in an Area of High Malaria Transmission. Front Immunol. 2017;8:1329. Epub 2017/11/04. doi: 10.3389/fimmu.2017.01329. PubMed PMID: 29097996; PubMed Central PMCID: PMCPMC5653696.

22. Jagannathan P, Eccles-James I, Bowen K, Nankya F, Auma A, Wamala S, et al. IFNgamma/IL-10 co-producing cells dominate the CD4 response to malaria in highly exposed children. PLoS Pathog. 2014;10(1):e1003864. Epub 2014/01/15. doi: 10.1371/journal.ppat.1003864. PubMed PMID: 24415936; PubMed Central PMCID: PMCPMC3887092.

23. Walther M, Jeffries D, Finney OC, Njie M, Ebonyi A, Deininger S, et al. Distinct roles for FOXP3 and FOXP3 CD4 T cells in regulating cellular immunity to uncomplicated and severe Plasmodium falciparum malaria. PLoS Pathog. 2009;5(4):e1000364. Epub 2009/04/04. doi: 10.1371/journal.ppat.1000364. PubMed PMID: 19343213; PubMed Central PMCID: PMCPMC2658808.

24. Perez-Mazliah D, Nguyen MP, Hosking C, McLaughlin S, Lewis MD, Tumwine I, et al. Follicular Helper T Cells are Essential for the Elimination of Plasmodium Infection. EBioMedicine. 2017;24:216–30. Epub 2017/09/11. doi: 10.1016/j.ebiom.2017.08.030. PubMed PMID: 28888925; PubMed Central PMCID: PMCPMC5652023.

25. Butler NS, Moebius J, Pewe LL, Traore B, Doumbo OK, Tygrett LT, et al. Therapeutic blockade of PD-L1 and LAG-3 rapidly clears established blood-stage Plasmodium infection. Nat Immunol. 2011;13(2):188–95. Epub 2011/12/14. doi: 10.1038/ni.2180. PubMed PMID: 22157630; PubMed Central PMCID: PMCPMC3262959.

26. Chan JA, Loughland JR, de Labastida Rivera F, SheelaNair A, Andrew DW, Dooley NL, et al. Th2-like T Follicular Helper Cells Promote Functional Antibody Production during Plasmodium falciparum Infection. Cell Rep Med. 2020;1(9):100157. Epub 2020/12/31. doi: 10.1016/j.xcrm.2020.100157. PubMed PMID: 33377128; PubMed Central PMCID: PMCPMC7762767.

27. Obeng-Adjei N, Portugal S, Tran TM, Yazew TB, Skinner J, Li S, et al. Circulating Th1-Cell-type Tfh Cells that Exhibit Impaired B Cell Help Are Preferentially Activated during Acute Malaria in Children. Cell Rep. 2015;13(2):425–39. Epub 2015/10/07. doi: 10.1016/j.celrep.2015.09.004. PubMed PMID: 26440897; PubMed Central PMCID: PMCPMC4607674.

28. Chan JA, Loughland JR, de la Parte L, Okano S, Ssewanyana I, Nalubega M, et al. Age-dependent changes in circulating Tfh cells influence development of functional malaria antibodies in children. Nat Commun. 2022;13(1):4159. Epub 2022/07/20. doi: 10.1038/s41467-022-31880-6. PubMed PMID: 35851033; PubMed Central PMCID: PMCPMC9293980.

29. Oyong DA, Loughland JR, Soon MSF, Chan JA, Andrew D, Wines BD, et al. Adults with Plasmodium falciparum malaria have higher magnitude and quality of circulating T-follicular helper cells compared to children. EBioMedicine. 2022;75:103784. Epub 2021/12/31. doi: 10.1016/j.ebiom.2021.103784. PubMed PMID: 34968760; PubMed Central PMCID: PMCPMC8718734.

30. Sedegah M, Sim BK, Mason C, Nutman T, Malik A, Roberts C, et al. Naturally acquired CD8+ cytotoxic T lymphocytes against the Plasmodium falciparum circumsporozoite protein. J Immunol. 1992;149(3):966–71. Epub 1992/08/01. PubMed PMID: 1634778.

31. Romero P, Maryanski JL, Corradin G, Nussenzweig RS, Nussenzweig V, Zavala F. Cloned cytotoxic T cells recognize an epitope in the circumsporozoite protein and protect against malaria. Nature. 1989;341(6240):323-6. Epub 1989/09/28. doi: 10.1038/341323a0. PubMed PMID: 2477703.

32. Epstein JE, Tewari K, Lyke KE, Sim BK, Billingsley PF, Laurens MB, et al. Live attenuated malaria vaccine designed to protect through hepatic CD8(+) T cell immunity. Science. 2011;334(6055):475-80. Epub 2011/09/10. doi: 10.1126/science.1211548. PubMed PMID: 21903775.

33. Van Braeckel-Budimir N, Harty JT. CD8 T-cell-mediated protection against liver-stage malaria: lessons from a mouse model. Front Microbiol. 2014;5:272. Epub 2014/06/18. doi: 10.3389/fmicb.2014.00272. PubMed PMID: 24936199; PubMed Central PMCID: PMCPMC4047659.

34. Weiss WR, Sedegah M, Beaudoin RL, Miller LH, Good MF. CD8+ T cells (cytotoxic/suppressors) are required for protection in mice immunized with malaria sporozoites. Proc Natl Acad Sci U S A. 1988;85(2):573–6. Epub 1988/01/01. doi: 10.1073/pnas.85.2.573. PubMed PMID: 2963334; PubMed Central PMCID: PMCPMC279593.

35. Schmidt NW, Butler NS, Badovinac VP, Harty JT. Extreme CD8 T cell requirements for anti-malarial liver-stage immunity following immunization with radiation attenuated sporozoites. PLoS Pathog. 2010;6(7):e1000998. Epub 2010/07/27. doi: 10.1371/journal.ppat.1000998. PubMed PMID: 20657824; PubMed Central PMCID: PMCPMC2904779.

36. Chakravarty S, Cockburn IA, Kuk S, Overstreet MG, Sacci JB, Zavala F. CD8+ T lymphocytes protective against malaria liver stages are primed in skin-draining lymph nodes. Nat Med. 2007;13(9):1035–41. Epub 2007/08/21. doi: 10.1038/nm1628. PubMed PMID: 17704784.

37. Radtke AJ, Kastenmuller W, Espinosa DA, Gerner MY, Tse SW, Sinnis P, et al. Lymph-node resident CD8alpha+ dendritic cells capture antigens from migratory malaria sporozoites and induce CD8+ T cell responses. PLoS Pathog. 2015;11(2):e1004637. Epub 2015/02/07. doi: 10.1371/journal.ppat.1004637. PubMed PMID: 25658939; PubMed Central PMCID: PMCPMC4450069.

38. Kurup SP, Anthony SM, Hancox LS, Vijay R, Pewe LL, Moioffer SJ, et al. Monocyte-Derived CD11c(+) Cells Acquire Plasmodium from Hepatocytes to Prime CD8 T Cell Immunity to Liver-Stage Malaria. Cell Host Microbe. 2019;25(4):565–77 e6. Epub 2019/03/25. doi: 10.1016/j.chom.2019.02.014. PubMed PMID: 30905437; PubMed Central PMCID: PMCPMC6459714.

39. Butler NS, Schmidt NW, Harty JT. Differential effector pathways regulate memory CD8 T cell immunity against Plasmodium berghei versus P. yoelii sporozoites. J Immunol. 2010;184(5):2528–38. Epub 2010/01/26. doi: 10.4049/jimmunol.0903529. PubMed PMID: 20097864; PubMed Central PMCID: PMCPMC2904689.

40. Renggli J, Hahne M, Matile H, Betschart B, Tschopp J, Corradin G. Elimination of P. berghei liver stages is independent of Fas (CD95/Apo-I) or perforin-mediated cytotoxicity. Parasite Immunol. 1997;19(3):145–8. Epub 1997/03/01. doi: 10.1046/j.1365-3024.1997.d01-190.x. PubMed PMID: 9106820.

41. Clark IA, Hunt NH, Butcher GA, Cowden WB. Inhibition of murine malaria (Plasmodium chabaudi) in vivo by recombinant interferon-gamma or tumor necrosis factor, and its enhancement by butylated hydroxyanisole. J Immunol. 1987;139(10):3493–6. Epub 1987/11/15. PubMed PMID: 3119710.

42. Holz LE, Prier JE, Freestone D, Steiner TM, English K, Johnson DN, et al. CD8(+) T Cell Activation Leads to Constitutive Formation of Liver Tissue-Resident Memory T Cells that Seed a Large and Flexible Niche in the Liver. Cell Rep. 2018;25(1):68–79 e4. Epub 2018/10/04. doi: 10.1016/j.celrep.2018.08.094. PubMed PMID: 30282039.

43. McNamara HA, Cai Y, Wagle MV, Sontani Y, Roots CM, Miosge LA, et al. Up-regulation of LFA-1 allows liver-resident memory T cells to patrol and remain in the hepatic sinusoids. Sci Immunol. 2017;2(9). Epub 2017/07/15. doi: 10.1126/sciimmunol.aaj1996. PubMed PMID: 28707003; PubMed Central PMCID: PMCPMC5505664.

44. de Menezes MN, Ge Z, Cozijnsen A, Gras S, Bertolino P, Caminschi I, et al. Long lived liver-resident memory T cells of biased specificities for abundant sporozoite antigens drive malaria protection by radiation-attenuated sporozoite vaccination. PLoS Pathog. 2025;21(5):e1012731. Epub 2025/05/27. doi: 10.1371/journal.ppat.1012731. PubMed PMID: 40424562.

45. Ganley M, Holz LE, Minnell JJ, de Menezes MN, Burn OK, Poa KCY, et al. mRNA vaccine against malaria tailored for liver-resident memory T cells. Nat Immunol. 2023;24(9):1487–98. Epub 2023/07/21. doi: 10.1038/s41590-023-01562-6. PubMed PMID: 37474653.

46. Kumar S, Miller LH. Cellular mechanisms in immunity to blood stage infection. Immunol Lett. 1990;25(1-3):109–14. Epub 1990/08/01. doi: 10.1016/0165-2478(90)90100-5. PubMed PMID: 1980907.

47. Junqueira C, Barbosa CRR, Costa PAC, Teixeira-Carvalho A, Castro G, Sen Santara S, et al. Cytotoxic CD8(+) T cells recognize and kill Plasmodium vivax-infected reticulocytes. Nat Med. 2018;24(9):1330–6. Epub 2018/07/25. doi: 10.1038/s41591-018-0117-4. PubMed PMID: 30038217; PubMed Central PMCID: PMCPMC6129205.

48. Claser C, Malleret B, Gun SY, Wong AY, Chang ZW, Teo P, et al. CD8+ T cells and IFN-gamma mediate the time-dependent accumulation of infected red blood cells in deep organs during experimental cerebral malaria. PLoS One. 2011;6(4):e18720. Epub 2011/04/16. doi: 10.1371/journal.pone.0018720. PubMed PMID: 21494565; PubMed Central PMCID: PMCPMC3073989 following conflicts: Laurent Renia is an academic editor of PLoS ONE.

49. Haque A, Best SE, Unosson K, Amante FH, de Labastida F, Anstey NM, et al. Granzyme B expression by CD8+ T cells is required for the development of experimental cerebral malaria. J Immunol. 2011;186(11):6148–56. Epub 2011/04/29. doi: 10.4049/jimmunol.1003955. PubMed PMID: 21525386.

50. Howland SW, Poh CM, Gun SY, Claser C, Malleret B, Shastri N, et al. Brain microvessel cross-presentation is a hallmark of experimental cerebral malaria. EMBO Mol Med. 2013;5(7):984–99. Epub 2013/05/18. doi: 10.1002/emmm.201202273. PubMed PMID: 23681698; PubMed Central PMCID: PMCPMC3721469.

51. Shafi AM, Vegvari A, Zubarev RA, Penha-Goncalves C. Brain endothelial cells exposure to malaria parasites links type I interferon signalling to antigen presentation, immunoproteasome activation, endothelium disruption, and cellular metabolism. Front Immunol. 2023;14:1149107. Epub 2023/03/31. doi: 10.3389/fimmu.2023.1149107. PubMed PMID: 36993973; PubMed Central PMCID: PMCPMC10042232.

52. Fernandes P, Howland SW, Heiss K, Hoffmann A, Hernandez-Castaneda MA, Obrova K, et al. A Plasmodium Cross-Stage Antigen Contributes to the Development of Experimental Cerebral Malaria. Front Immunol. 2018;9:1875. Epub 2018/08/30. doi: 10.3389/fimmu.2018.01875. PubMed PMID: 30154793; PubMed Central PMCID: PMCPMC6102508.

53. Ng SS, De Labastida Rivera F, Yan J, Corvino D, Das I, Zhang P, et al. The NK cell granule protein NKG7 regulates cytotoxic granule exocytosis and inflammation. Nat Immunol. 2020;21(10):1205–18. Epub 2020/08/26. doi: 10.1038/s41590-020-0758-6. PubMed PMID: 32839608; PubMed Central PMCID: PMCPMC7965849.

54. Houchins JP, Yabe T, McSherry C, Miyokawa N, Bach FH. Isolation and characterization of NK cell or NK/T cell-specific cDNA clones. J Mol Cell Immunol. 1990;4(6):295–304; discussion 5-6. Epub 1990/01/01. PubMed PMID: 2080984.

55. Mori S, Jewett A, Cavalcanti M, Murakami-Mori K, Nakamura S, Bonavida B. Differential regulation of human NK cell-associated gene expression following activation by IL-2, IFN-alpha and PMA/ionomycin. Int J Oncol. 1998;12(5):1165–70. Epub 1998/06/06. doi: 10.3892/ijo.12.5.1165. PubMed PMID: 9538144.

56. Turman MA, Yabe T, McSherry C, Bach FH, Houchins JP. Characterization of a novel gene (NKG7) on human chromosome 19 that is expressed in natural killer cells and T cells. Hum Immunol. 1993;36(1):34–40. Epub 1993/01/01. doi: 10.1016/0198-8859(93)90006-m. PubMed PMID: 8458737.

57. Kawakami A, Tian Q, Streuli M, Poe M, Edelhoff S, Disteche CM, et al. Intron-exon organization and chromosomal localization of the human TIA-1 gene. J Immunol. 1994;152(10):4937–45. Epub 1994/05/15. PubMed PMID: 8176212.

58. Medley QG, Kedersha N, O’Brien S, Tian Q, Schlossman SF, Streuli M, et al. Characterization of GMP-17, a granule membrane protein that moves to the plasma membrane of natural killer cells following target cell recognition. Proc Natl Acad Sci U S A. 1996;93(2):685–9. Epub 1996/01/23. doi: 10.1073/pnas.93.2.685. PubMed PMID: 8570616; PubMed Central PMCID: PMCPMC40113.

59. Uhlen M, Fagerberg L, Hallstrom BM, Lindskog C, Oksvold P, Mardinoglu A, et al. Proteomics. Tissue-based map of the human proteome. Science. 2015;347(6220):1260419. Epub 2015/01/24. doi: 10.1126/science.1260419. PubMed PMID: 25613900.

60. Zaunders JJ, Dyer WB, Wang B, Munier ML, Miranda-Saksena M, Newton R, et al. Identification of circulating antigen-specific CD4+ T lymphocytes with a CCR5+, cytotoxic phenotype in an HIV-1 long-term nonprogressor and in CMV infection. Blood. 2004;103(6):2238–47. Epub 2003/12/03. doi: 10.1182/blood-2003-08-2765. PubMed PMID: 14645006.

61. Pachnio A, Ciaurriz M, Begum J, Lal N, Zuo J, Beggs A, et al. Cytomegalovirus Infection Leads to Development of High Frequencies of Cytotoxic Virus-Specific CD4+ T Cells Targeted to Vascular Endothelium. PLoS Pathog. 2016;12(9):e1005832. Epub 2016/09/09. doi: 10.1371/journal.ppat.1005832. PubMed PMID: 27606804; PubMed Central PMCID: PMCPMC5015996.

62. Thelen B, Schipperges V, Knorlein P, Hummel JF, Arnold F, Kupferschmid L, et al. Eomes is sufficient to regulate IL-10 expression and cytotoxic effector molecules in murine CD4(+) T cells. Front Immunol. 2023;14:1058267. Epub 2023/02/10. doi: 10.3389/fimmu.2023.1058267. PubMed PMID: 36756120; PubMed Central PMCID: PMCPMC9901365.

63. Fairfax BP, Taylor CA, Watson RA, Nassiri I, Danielli S, Fang H, et al. Peripheral CD8(+) T cell characteristics associated with durable responses to immune checkpoint blockade in patients with metastatic melanoma. Nat Med. 2020;26(2):193–9. Epub 2020/02/12. doi: 10.1038/s41591-019-0734-6. PubMed PMID: 32042196; PubMed Central PMCID: PMCPMC7611047.

64. Xu JL. Wilms Tumor 1-Associated Protein Expression Is Linked to a T-Cell-Inflamed Phenotype in Pancreatic Cancer. Dig Dis Sci. 2023;68(3):831–40. Epub 2022/07/22. doi: 10.1007/s10620-022-07620-7. PubMed PMID: 35859262.

65. Jenner RG, Townsend MJ, Jackson I, Sun K, Bouwman RD, Young RA, et al. The transcription factors T-bet and GATA-3 control alternative pathways of T-cell differentiation through a shared set of target genes. Proc Natl Acad Sci U S A. 2009;106(42):17876–81. Epub 2009/10/07. doi: 10.1073/pnas.0909357106. PubMed PMID: 19805038; PubMed Central PMCID: PMCPMC2764903 and holds equity in the Bristol Myers Squibb Corporation.

66. Hahtola S, Tuomela S, Elo L, Hakkinen T, Karenko L, Nedoszytko B, et al. Th1 response and cytotoxicity genes are down-regulated in cutaneous T-cell lymphoma. Clin Cancer Res. 2006;12(16):4812–21. Epub 2006/08/18. doi: 10.1158/1078-0432.CCR-06-0532. PubMed PMID: 16914566.

67. Koch MA, Thomas KR, Perdue NR, Smigiel KS, Srivastava S, Campbell DJ. T-bet(+) Treg cells undergo abortive Th1 cell differentiation due to impaired expression of IL-12 receptor beta2. Immunity. 2012;37(3):501–10. Epub 2012/09/11. doi: 10.1016/j.immuni.2012.05.031. PubMed PMID: 22960221; PubMed Central PMCID: PMCPMC3501343.

68. Zhang J-Y, Wang X-M, Xing X, Xu Z, Zhang C, Song J-W, et al. Single-cell landscape of immunological responses in patients with COVID-19. Nature Immunology. 2020;21(9):1107–18. doi: 10.1038/s41590-020-0762-x.

69. Yan M, Cao H, Tao K, Xiao B, Chu Y, Ma D, et al. HDACs alters negatively to the tumor immune microenvironment in gynecologic cancers. Gene. 2023;885:147704. Epub 2023/08/13. doi: 10.1016/j.gene.2023.147704. PubMed PMID: 37572797.

70. Li DF, Tulahong A, Uddin MN, Zhao H, Zhang H. Meta-analysis identifying epithelial-derived transcriptomes predicts poor clinical outcome and immune infiltrations in ovarian cancer. Math Biosci Eng. 2021;18(5):6527–51. Epub 2021/09/15. doi: 10.3934/mbe.2021324. PubMed PMID: 34517544.

71. Imeri J, Desterke C, Marcoux P, Chaker D, Oudrhiri N, Fund X, et al. Case report: Long-term voluntary Tyrosine Kinase Inhibitor (TKI) discontinuation in chronic myeloid leukemia (CML): Molecular evidence of an immune surveillance. Front Oncol. 2023;13:1117781. Epub 2023/04/04. doi: 10.3389/fonc.2023.1117781. PubMed PMID: 37007090; PubMed Central PMCID: PMCPMC10062417.

72. Xue S, Su XM, Ke LN, Huang YG. CXCL9 correlates with antitumor immunity and is predictive of a favorable prognosis in uterine corpus endometrial carcinoma. Front Oncol. 2023;13:1077780. Epub 2023/02/28. doi: 10.3389/fonc.2023.1077780. PubMed PMID: 36845675; PubMed Central PMCID: PMCPMC9945585.

73. Casarrubios M, Provencio M, Nadal E, Insa A, Del Rosario Garcia-Campelo M, Lazaro-Quintela M, et al. Tumor microenvironment gene expression profiles associated to complete pathological response and disease progression in resectable NSCLC patients treated with neoadjuvant chemoimmunotherapy. J Immunother Cancer. 2022;10(9). Epub 2022/09/29. doi: 10.1136/jitc-2022-005320. PubMed PMID: 36171009; PubMed Central PMCID: PMCPMC9528578.

74. Myles A, Tuteja A, Aggarwal A. Synovial fluid mononuclear cell gene expression profiling suggests dysregulation of innate immune genes in enthesitis-related arthritis patients. Rheumatology (Oxford). 2012;51(10):1785–9. Epub 2012/07/06. doi: 10.1093/rheumatology/kes151. PubMed PMID: 22763987.

75. Lit L, Gilbert DL, Walker W, Sharp FR. A subgroup of Tourette’s patients overexpress specific natural killer cell genes in blood: a preliminary report. Am J Med Genet B Neuropsychiatr Genet. 2007;144B(7):958–63. Epub 2007/05/16. doi: 10.1002/ajmg.b.30550. PubMed PMID: 17503477.

76. Lit L, Enstrom A, Sharp FR, Gilbert DL. Age-related gene expression in Tourette syndrome. J Psychiatr Res. 2009;43(3):319–30. Epub 2008/05/20. doi: 10.1016/j.jpsychires.2008.03.012. PubMed PMID: 18485367; PubMed Central PMCID: PMCPMC2662336.

77. Li XY, Corvino D, Nowlan B, Aguilera AR, Ng SS, Braun M, et al. NKG7 Is Required for Optimal Antitumor T-cell Immunity. Cancer Immunol Res. 2022;10(2):154–61. Epub 2022/01/12. doi: 10.1158/2326-6066.CIR-20-0649. PubMed PMID: 35013002.

78. Fernandez-Ruiz D, Lau LS, Ghazanfari N, Jones CM, Ng WY, Davey GM, et al. Development of a Novel CD4(+) TCR Transgenic Line That Reveals a Dominant Role for CD8(+) Dendritic Cells and CD40 Signaling in the Generation of Helper and CTL Responses to Blood-Stage Malaria. J Immunol. 2017;199(12):4165–79. Epub 2017/11/01. doi: 10.4049/jimmunol.1700186. PubMed PMID: 29084838; PubMed Central PMCID: PMCPMC5713497.

79. Engwerda CR, Minigo G, Amante FH, McCarthy JS. Experimentally induced blood stage malaria infection as a tool for clinical research. Trends Parasitol. 2012;28(11):515–21. Epub 2012/10/09. doi: 10.1016/j.pt.2012.09.001. PubMed PMID: 23041118.

80. Abd-Rahman AN, Kaschek D, Kümmel A, Webster R, Potter AJ, Odedra A, et al. Characterizing the pharmacological interaction of the antimalarial combination artefenomel-piperaquine in healthy volunteers with induced blood-stage Plasmodium falciparum to predict efficacy in patients with malaria. BMC Med. 2024;22(1):563. Epub 2024/11/29. doi: 10.1186/s12916-024-03787-0. PubMed PMID: 39609822; PubMed Central PMCID: PMCPMC11603672.

81. Choi SJ, Koh J-Y, Rha M-S, Seo I-H, Lee H, Jeong S, et al. KIR+CD8+ and NKG2A+CD8+ T cells are distinct innate-like populations in humans. Cell Reports. 2023;42(3):112236. doi: 10.1016/j.celrep.2023.112236.

82. Utzschneider DT, Charmoy M, Chennupati V, Pousse L, Ferreira DP, Calderon-Copete S, et al. T Cell Factor 1-Expressing Memory-like CD8+ T Cells Sustain the Immune Response to Chronic Viral Infections. Immunity. 2016;45(2):415–27. doi: 10.1016/j.immuni.2016.07.021.

83. Kurup SP, Butler NS, Harty JT. T cell-mediated immunity to malaria. Nat Rev Immunol. 2019;19(7):457–71. Epub 2019/04/04. doi: 10.1038/s41577-019-0158-z. PubMed PMID: 30940932; PubMed Central PMCID: PMCPMC6599480.

84. Hassert M, Arumugam S, Harty JT. Memory CD8+ T cell-mediated protection against liver-stage malaria. Immunol Rev. 2023;316(1):84–103. Epub 2023/04/05. doi: 10.1111/imr.13202. PubMed PMID: 37014087.

85. Okamura T, Hamaguchi M, Tominaga H, Kitagawa N, Hashimoto Y, Majima S, et al. Characterization of Peripheral Blood TCR in Patients with Type 1 Diabetes Mellitus by BD Rhapsody(TM) VDJ CDR3 Assay. Cells. 2022;11(10). Epub 2022/05/29. doi: 10.3390/cells11101623. PubMed PMID: 35626661; PubMed Central PMCID: PMCPMC9139223.

86. Alber S, Kumar S, Liu J, Huang ZM, Paez D, Hong J, et al. Single Cell Transcriptome and Surface Epitope Analysis of Ankylosing Spondylitis Facilitates Disease Classification by Machine Learning. Front Immunol. 2022;13:838636. Epub 2022/06/01. doi: 10.3389/fimmu.2022.838636. PubMed PMID: 35634297; PubMed Central PMCID: PMCPMC9135966.

87. Zheng M, Zhou W, Huang C, Hu Z, Zhang B, Lu Q, et al. A single-cell map of peripheral alterations after FMT treatment in patients with systemic lupus erythematosus. J Autoimmun. 2023;135:102989. Epub 2023/01/08. doi: 10.1016/j.jaut.2022.102989. PubMed PMID: 36610264.

88. Lagumdzic E, Pernold CPS, Ertl R, Palmieri N, Stadler M, Sawyer S, et al. Gene expression of peripheral blood mononuclear cells and CD8(+) T cells from gilts after PRRSV infection. Front Immunol. 2023;14:1159970. Epub 2023/07/06. doi: 10.3389/fimmu.2023.1159970. PubMed PMID: 37409113; PubMed Central PMCID: PMCPMC10318438.

89. Wen T, Barham W, Li Y, Zhang H, Gicobi JK, Hirdler JB, et al. NKG7 Is a T-cell-Intrinsic Therapeutic Target for Improving Antitumor Cytotoxicity and Cancer Immunotherapy. Cancer Immunol Res. 2022;10(2):162–81. Epub 2021/12/17. doi: 10.1158/2326-6066.CIR-21-0539. PubMed PMID: 34911739; PubMed Central PMCID: PMCPMC8816890.

90. Morikawa Y, Murakami M, Kondo H, Nemoto N, Iwabuchi K, Eshima K. Natural Killer Cell Group 7 Sequence in Cytotoxic Cells Optimizes Exocytosis of Lytic Granules Essential for the Perforin-Dependent, but Not Fas Ligand-Dependent, Cytolytic Pathway. Immunohorizons. 2021;5(4):234–45. Epub 2021/04/30. doi: 10.4049/immunohorizons.2100029. PubMed PMID: 33911019.

91. Lelliott EJ, Ramsbottom KM, Dowling MR, Shembrey C, Noori T, Kearney CJ, et al. NKG7 Enhances CD8+ T Cell Synapse Efficiency to Limit Inflammation. Front Immunol. 2022;13:931630. Epub 2022/07/26. doi: 10.3389/fimmu.2022.931630. PubMed PMID: 35874669; PubMed Central PMCID: PMCPMC9299089.

92. Kong G, Song Y, Yan Y, Calderazzo SM, Saddala MS, De Labastida Rivera F, et al. Clonally expanded, targetable, natural killer-like NKG7 T cells seed the aged spinal cord to disrupt myeloid-dependent wound healing. Neuron. 2025;113(5):684–700.e8. Epub 2025/01/15. doi: 10.1016/j.neuron.2024.12.012. PubMed PMID: 39809279.

93. Chen Y, Wang M, Huang S, Han L, Cai Y, Xu X, et al. Ectopic expression of NKG7 enhances CAR-T function and improves the therapeutic efficacy in liquid and solid tumors. Pharmacological Research. 2024;210:107506. doi: 10.1016/j.phrs.2024.107506.

94. Gbedande K, Carpio VH, Stephens R. Using two phases of the CD4 T cell response to blood-stage murine malaria to understand regulation of systemic immunity and placental pathology in Plasmodium falciparum infection. Immunol Rev. 2020;293(1):88–114. Epub 2020/01/07. doi: 10.1111/imr.12835. PubMed PMID: 31903675; PubMed Central PMCID: PMCPMC7540220.

95. Freuchet A, Roy P, Armstrong SS, Oliaeimotlagh M, Kumar S, Orecchioni M, et al. Identification of human exT(reg) cells as CD16(+)CD56(+) cytotoxic CD4(+) T cells. Nat Immunol. 2023. Epub 2023/08/11. doi: 10.1038/s41590-023-01589-9. PubMed PMID: 37563308.

96. Lund R, Ahlfors H, Kainonen E, Lahesmaa AM, Dixon C, Lahesmaa R. Identification of genes involved in the initiation of human Th1 or Th2 cell commitment. Eur J Immunol. 2005;35(11):3307–19. Epub 2005/10/13. doi: 10.1002/eji.200526079. PubMed PMID: 16220538.

97. Iwata S, Mikami Y, Sun HW, Brooks SR, Jankovic D, Hirahara K, et al. The Transcription Factor T-bet Limits Amplification of Type I IFN Transcriptome and Circuitry in T Helper 1 Cells. Immunity. 2017;46(6):983–91 e4. Epub 2017/06/18. doi: 10.1016/j.immuni.2017.05.005. PubMed PMID: 28623086; PubMed Central PMCID: PMCPMC5523825.

98. Gokmen MR, Dong R, Kanhere A, Powell N, Perucha E, Jackson I, et al. Genome-wide regulatory analysis reveals that T-bet controls Th17 lineage differentiation through direct suppression of IRF4. J Immunol. 2013;191(12):5925–32. Epub 2013/11/20. doi: 10.4049/jimmunol.1202254. PubMed PMID: 24249732; PubMed Central PMCID: PMCPMC3858236.

99. Ashokkumar C, Ningappa M, Ranganathan S, Higgs BW, Sun Q, Schmitt L, et al. Increased expression of peripheral blood leukocyte genes implicate CD14+ tissue macrophages in cellular intestine allograft rejection. Am J Pathol. 2011;179(4):1929–38. Epub 2011/08/23. doi: 10.1016/j.ajpath.2011.06.040. PubMed PMID: 21854741; PubMed Central PMCID: PMCPMC3181350.

100. Khatri P, Roedder S, Kimura N, De Vusser K, Morgan AA, Gong Y, et al. A common rejection module (CRM) for acute rejection across multiple organs identifies novel therapeutics for organ transplantation. J Exp Med. 2013;210(11):2205–21. Epub 2013/10/16. doi: 10.1084/jem.20122709. PubMed PMID: 24127489; PubMed Central PMCID: PMCPMC3804941.

101. Sigdel TK, Bestard O, Tran TQ, Hsieh SC, Roedder S, Damm I, et al. A Computational Gene Expression Score for Predicting Immune Injury in Renal Allografts. PLoS One. 2015;10(9):e0138133. Epub 2015/09/15. doi: 10.1371/journal.pone.0138133. PubMed PMID: 26367000; PubMed Central PMCID: PMCPMC4569485.

102. Pidala J, Bloom GC, Eschrich S, Sarwal M, Enkemann S, Betts BC, et al. Tolerance associated gene expression following allogeneic hematopoietic cell transplantation. PLoS One. 2015;10(3):e0117001. Epub 2015/03/17. doi: 10.1371/journal.pone.0117001. PubMed PMID: 25774806; PubMed Central PMCID: PMCPMC4361657.

103. T GM, Gauthier CD, Murphy L, Lanser TB, Paul A, Matos KTF, et al. Nasal administration of anti-CD3 mAb (Foralumab) downregulates NKG7 and increases TGFB1 and GIMAP7 expression in T cells in subjects with COVID-19. Proc Natl Acad Sci U S A. 2023;120(11):e2220272120. Epub 2023/03/08. doi: 10.1073/pnas.2220272120. PubMed PMID: 36881624; PubMed Central PMCID: PMCPMC10243127.

104. M MN, Newell F, Aoude LG, Bonazzi VF, Patel K, Lampe G, et al. Multi-omic features of oesophageal adenocarcinoma in patients treated with preoperative neoadjuvant therapy. Nat Commun. 2023;14(1):3155. Epub 2023/06/01. doi: 10.1038/s41467-023-38891-x. PubMed PMID: 37258531; PubMed Central PMCID: PMCPMC10232490 other authors have no competing interests.

105. Yu W, Wang S, Rong Q, Ajayi OE, Hu K, Wu Q. Profiling the Tumor-Infiltrating Lymphocytes in Gastric Cancer Reveals Its Implication in the Prognosis. Genes (Basel). 2022;13(6). Epub 2022/06/25. doi: 10.3390/genes13061017. PubMed PMID: 35741779; PubMed Central PMCID: PMCPMC9222794.

106. Lau LS, Fernandez-Ruiz D, Mollard V, Sturm A, Neller MA, Cozijnsen A, et al. CD8+ T cells from a novel T cell receptor transgenic mouse induce liver-stage immunity that can be boosted by blood-stage infection in rodent malaria. PLoS Pathog. 2014;10(5):e1004135. Epub 2014/05/24. doi: 10.1371/journal.ppat.1004135. PubMed PMID: 24854165; PubMed Central PMCID: PMCPMC4031232.

107. Barber BE, Grigg MJ, William T, Piera KA, Boyle MJ, Yeo TW, et al. Effects of Aging on Parasite Biomass, Inflammation, Endothelial Activation, Microvascular Dysfunction and Disease Severity in Plasmodium knowlesi and Plasmodium falciparum Malaria. J Infect Dis. 2017;215(12):1908–17. Epub 2017/09/03. doi: 10.1093/infdis/jix193. PubMed PMID: 28863470; PubMed Central PMCID: PMCPMC8453637.

108. Barber BE, William T, Grigg MJ, Parameswaran U, Piera KA, Price RN, et al. Parasite biomass-related inflammation, endothelial activation, microvascular dysfunction and disease severity in vivax malaria. PLoS Pathog. 2015;11(1):e1004558. Epub 2015/01/09. doi: 10.1371/journal.ppat.1004558. PubMed PMID: 25569250; PubMed Central PMCID: PMCPMC4287532.

109. Marcus E. A STAR Is Born. Cell. 2016;166(5):1059–60. doi: 10.1016/j.cell.2016.08.021.

110. Franke-Fayard B, Janse CJ, Cunha-Rodrigues M, Ramesar J, Buscher P, Que I, et al. Murine malaria parasite sequestration: CD36 is the major receptor, but cerebral pathology is unlinked to sequestration. Proc Natl Acad Sci U S A. 2005;102(32):11468–73. Epub 2005/07/30. doi: 10.1073/pnas.0503386102. PubMed PMID: 16051702; PubMed Central PMCID: PMCPMC1183563.

